# Engineering high-titer lentiviral vectors for robust expression of RNA-based gene circuits

**DOI:** 10.64898/2026.05.12.724401

**Authors:** Kasey S. Love, Brittany A. Lende-Dorn, Kate E. Galloway

**Affiliations:** Department of Biological Engineering, MIT, 25 Ames St., Cambridge, MA 02139, USA; Department of Chemical Engineering, MIT, 25 Ames St., Cambridge, MA 02139, USA; Broad Institute of MIT and Harvard, 415 Main St., Cambridge, MA 02142, USA; Koch Institute for Integrative Cancer Research, 500 Main St., Cambridge, MA 02139, USA

## Abstract

Lentiviral vectors enable efficient delivery of genetic cargoes for gene and cell therapies. With their ∼10-kb packaging limit, lentiviral vectors can encode multiple transcription units, supporting delivery of compact gene circuits. RNA-based devices offer highly compact control including ligand-responsive induction and closed-loop regulation. However, RNA devices such as ribozymes and splicing switches may interfere with vector production via activity on the single-stranded RNA genome. Here, we examine the impact of gene syntax and genetic parts to define design strategies for two-gene vectors encoding RNA devices. We find that titer decreases with genetic parts that interfere with transcription or processing of the viral transcript during production. Compared to initial vectors, our best-performing design boosts titer more than 30-fold, enabling fine-scale tuning of expression to optimize cell-fate conversion within a nonmonotonic landscape. Together, this work illuminates principles for constructing two-gene lentiviral vectors with both high titer and robust expression, enhancing efficacy for downstream applications.

## Introduction

Lentiviral vectors are effective delivery methods for *ex vivo* engineering of cell therapies and *in vivo* gene therapies. Eleven lentiviral-mediated therapies are currently approved by the United States Food and Drug Administration^1^, and hundreds more have been registered in clinical trials^2^. These vectors have broad or programmable tropism and can transduce both dividing and non-dividing cells, enabling targeting to therapeutic cell types^2–6^. Additionally, lentiviral vectors offer stable, long-term expression, minimizing required dosages and enhancing efficacy^5^. Furthermore, the relatively large cargo capacity (∼10 kilobases) of these vectors allows for inclusion of multiple transcriptional units to deliver several cargoes or gene circuits^2,7,8^. Gene circuits afford sophisticated control, including autonomous regulation and response to exogenous inputs, that can contribute to greater safety and efficacy of therapeutic vectors.

Composing gene circuits with RNA devices is particularly advantageous, as these elements are highly compact, portable, and programmable^9^. RNA devices like microRNAs and ribozyme switches are typically no more than a few hundred base pairs, contributing a small footprint in size-limited delivery vectors. As these elements may not require exogenous protein components to function, they minimize immunogenicity and toxicity when expressed in target cells. RNA components may be sequence-programmable to interface with or act orthogonally to native networks, and ligand-responsive aptamers can be composed with functional domains via high-throughput design techniques^10–12^. RNA-based circuits with cell type-specificity^13,14^ or small-molecule control^15–23^ are increasingly being applied in therapeutic contexts. However, RNA-based regulatory elements that include splice sites, strong secondary or tertiary structures, or sequences targeted by machinery in producer cells may disrupt production of viral vectors used for delivery. Thus, there is an opportunity to define principles for designing lentiviral vectors containing these elements to unlock their clinical potential.

Effective lentiviral vectors require both efficient production and robust expression of their cargo^24,25^. If viral titer is low or the cargo expresses poorly, a larger dose of the vector is required to elicit the desired expression level in transduced target cells, limiting vector scale-up (Fig. **1A**).

**Figure 1.**
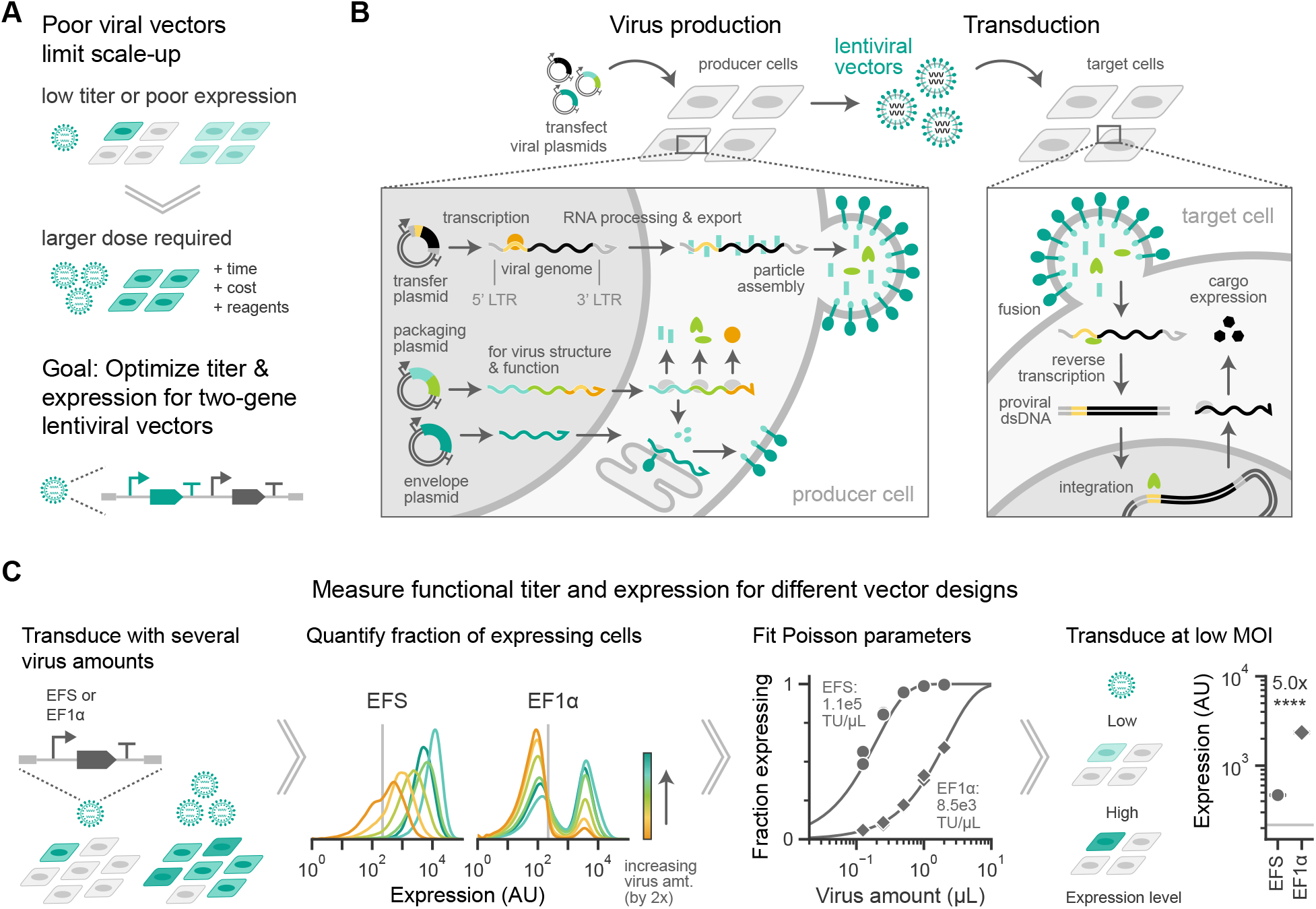
Promoter identity sets tradeoffs between viral titer and expression level of single genes. A. Viral vectors with low titer or poor expression require larger doses to elicit the desired behavior in target cells. Optimized vectors can improve production scale-up. B. Lentiviral vectors are packaged in producer cell lines transfected with a transfer plasmid to express the viral genome. The viral genome is a single-stranded RNA species spanning from the 5’ long terminal repeat sequence (LTR) to the 3’ LTR that contains the transgenic cargo along with other viral sequences that facilitate packaging. This transcript is exported from the nucleus and packaged with viral proteins into particles that bud from the membrane of producer cells. The viral particles are collected and then applied to target cells. During transduction, the envelope protein mediates fusion of the particle with the target cell membrane, releasing the particle contents into the cytoplasm. The viral genome is reverse transcribed into proviral double-stranded DNA (dsDNA), which is randomly integrated into the genome of the target cell. The target cell then expresses the transgenic cargo. C. Workflow to measure functional titer and expression for various vector designs. Single-gene vectors containing a fluorescent protein expressed by either the human elongation factor 1 alpha (EF1α) or EF1α short (EFS) promoter were transduced at varying amounts into HEK293T cells. **Left:** Expression of the single-gene vectors measured via fluorescence in arbitrary units (AU, log scale) from a flow cytometer. Light gray vertical line depicts the expression gate, used to calculate the fraction of expressing cells in each condition. Distributions show one representative batch of virus (one biological replicate). **Middle:** For the same vectors, fraction expressing as a function of virus volume was fit to a Poisson distribution to obtain titer in transducing units (TU) per µL virus. Points represent technical replicates for a representative batch of virus, and lines depict the Poisson curve fits. The computed titer is annotated on the plot, and virus volume is plotted on a log scale. **Right:** To quantify cargo expression, HEK293T cells were transduced at a low multiplicity of infection (MOI). Expression is the geometric mean of expressing cells (AU, log scale). Solid light gray line depicts the expression gate. Points represent means ±standard error for *n* = 3 biological replicates. Statistic is a two-sided Student’s t-test, **** *p <* 0.0001. Annotation shows fold change between indicated points.

Effective vector design may be able to optimize both titer and expression simultaneously, or designs may illuminate tradeoffs for application-specific decisions. These design choices may impact production and expression at several steps during lentiviral vector manufacturing and delivery^26^. Correct processing and packaging of the single-stranded RNA (ssRNA) viral genome in producer cells is essential for reverse transcription, integration, and expression of the desired cargo in transduced target cells (Fig. **1B**). Cargo sequences that disrupt any of these steps will reduce production and packaging of full-length genomes^27^. For instance, early termination during transcription of the viral genome in producer cells will generate transcripts that lack components required for integration of the proviral DNA in target cells^28–30^. Cryptic splicing between viral sequences and transgenic elements may produce transcripts that are packaged less efficiently or that integrate an unwanted, modified sequence in target cells^31–35^. Potential integration of unintended sequences is particularly problematic for *in vivo* gene therapy, as aberrant sequences cannot be easily removed. Alternatively, encoding disruptive sequences on the antisense strand of the cargo may limit unintended activity of strand-specific RNA elements on the ssRNA viral genome^31,33^. However, inverting an entire transcriptional unit may interfere with virus production. Strong expression of transcripts that are antisense to the primary viral transcript may inhibit transcription and processing of the desired viral genome or disrupt packaging into viral particles via formation of double-stranded RNA species^36–39^.

To support transgene expression in target cells, vector design must balance tradeoffs between viral titer and expression after integration. For vectors with two transcriptional units, expression is affected by gene syntax, the order and orientation of adjacent genes. We previously demonstrated that divergent syntax can amplify expression from both genes, while tandem syntax can reduce expression of the downstream gene due to DNA supercoiling-mediated feedback^40^. Additionally, readthrough transcription from the upstream gene in tandem syntax can reduce expression of one or both genes^24,41^. On the other hand, inclusion of a polyadenylation signal (PAS) on the upstream gene can mitigate readthrough^42,43^ and boost expression levels^44^. As expression of virally integrated transgenes can silence over time, some applications may require inclusion of sequences that help resist silencing^45–47^, even if these reduce viral titer. Thus, identifying tradeoffs in titer and expression is important for negotiating competing design objectives in downstream applications.

Here, we aimed to uncover how design choices affect both titer and expression. In order to optimize these properties, we systematically varied genetic elements and gene syntax in lentiviral vectors containing two transcriptional units. By quantifying both titer and expression, we identified vector designs that function well across cargoes and support expression of RNA devices in gene circuits, including a microRNA-based dosage controller, a ligand-responsive ribozyme switch, and a ligand-responsive splicing switch. Our best-performing vector design encodes the RNA devices under an inducible promoter antisense to the viral genome with divergent syntax, increasing titer more than 30-fold and maintaining functionality compared to alternatives arranged in tandem. This design enables fine-scale titration of functional genes in primary cells. By tuning expression of a biphasic cell-fate regulator, we identified the expression level at which we could optimize cell-fate conversion in primary mouse cells, highlighting the utility of our design principles in downstream applications.

## Results

### Promoter identity sets tradeoffs between viral titer and expression level of single genes

We set out to understand design rules for composing two transcriptional units on a single lentiviral vector, beginning with vectors containing a single transcriptional unit to establish a baseline. To quantify the effects of design choices on vector production and expression, we measured both functional titer and expression after single-copy integration in target cells (Fig. **1C**). Specifically, following standard virus production in HEK293T cells, we transduced fresh HEK293T cells with varying amounts of concentrated virus and computed the fraction of expressing cells three days later using flow cytometry. Assuming a Poisson process for transduction events, we fit the data to calculate titer in transducing units per volume of concentrated virus. We chose this functional metric for titer to capture both the particle formation and integration processes, as these together are relevant for downstream applications. Then, using these titer values, we transduced new HEK293T cells at a standard multiplicity of infection (MOI) of 0.3 to ensure single-copy integration and quantified expression level in transduced cells. Using fluorescent reporters, we could compare both viral titer and cargo expression across vector designs.

Promoter choice represents an important design consideration for specifying gene expression at the RNA and protein levels^48^. We first compared two commonly used promoters, measuring their effects on titer and expression. For single-gene vectors expressing a fluorescent protein from the human elongation factor 1 alpha (EF1α) promoter or a shorter, intronless variant (EF1α short, EFS), we observed a more than 10-fold difference in functional titer between the designs (Fig. **1C**). Potentially, the intron sequence associated with EF1α interferes with processing of the viral genome. Despite its lower titer, the vector encoding EF1α led to five-fold higher expression compared to the EFS design at the same MOI (Fig. **1C**), reflecting previously reported differences in expression strength between these two promoters^48^. Thus, we find that promoter identity can set tradeoffs between viral titer and expression level of single genes.

### Promoter strength and gene syntax define the tradeoffs in two-gene vectors

For two-gene vectors, gene syntax and genetic parts affect the RNA species generated in both producer and target cells. The orientation of cargo genes relative to the viral genome affects transcription and RNA processing during virus production (Fig. **2A**). In producer cells, a promoter on the transfer plasmid upstream of the 5’ LTR initiates transcription of the viral genome, which can be packaged into productive particles^49^. However, genetic elements on the sense strand can result in splicing, truncation, or other aberrant processing of the viral transcript, decreasing production of particles carrying the correct sequence^28–35^. In addition to the full-length, ssRNA viral genome (5’ LTR to 3’ LTR), promoters within the cargo of the vector may generate other transcripts. Antisense transcripts can interfere with transcription or processing of the viral genome or form double-stranded RNA species that inhibit packaging^36–38^. For two-gene vectors, the orientation and order of cargo genes (i.e., gene syntax) affects expression after integration (Fig. **2B**). In target cells, genes with tandem syntax may exhibit supercoiling-mediated upstream dominance^40^ or readthrough transcription^24,42,43^, decreasing expression of the downstream gene. In contrast, divergent syntax may amplify expression of both genes through biophysical coupling^40^. For both syntaxes, choice of genetic elements alters transcript processing, stability, and translation to impact expression^42,43,48^. Given these diverse processes, we sought to identify how gene syntax and components together influence both vector production and expression.

**Figure 2.**
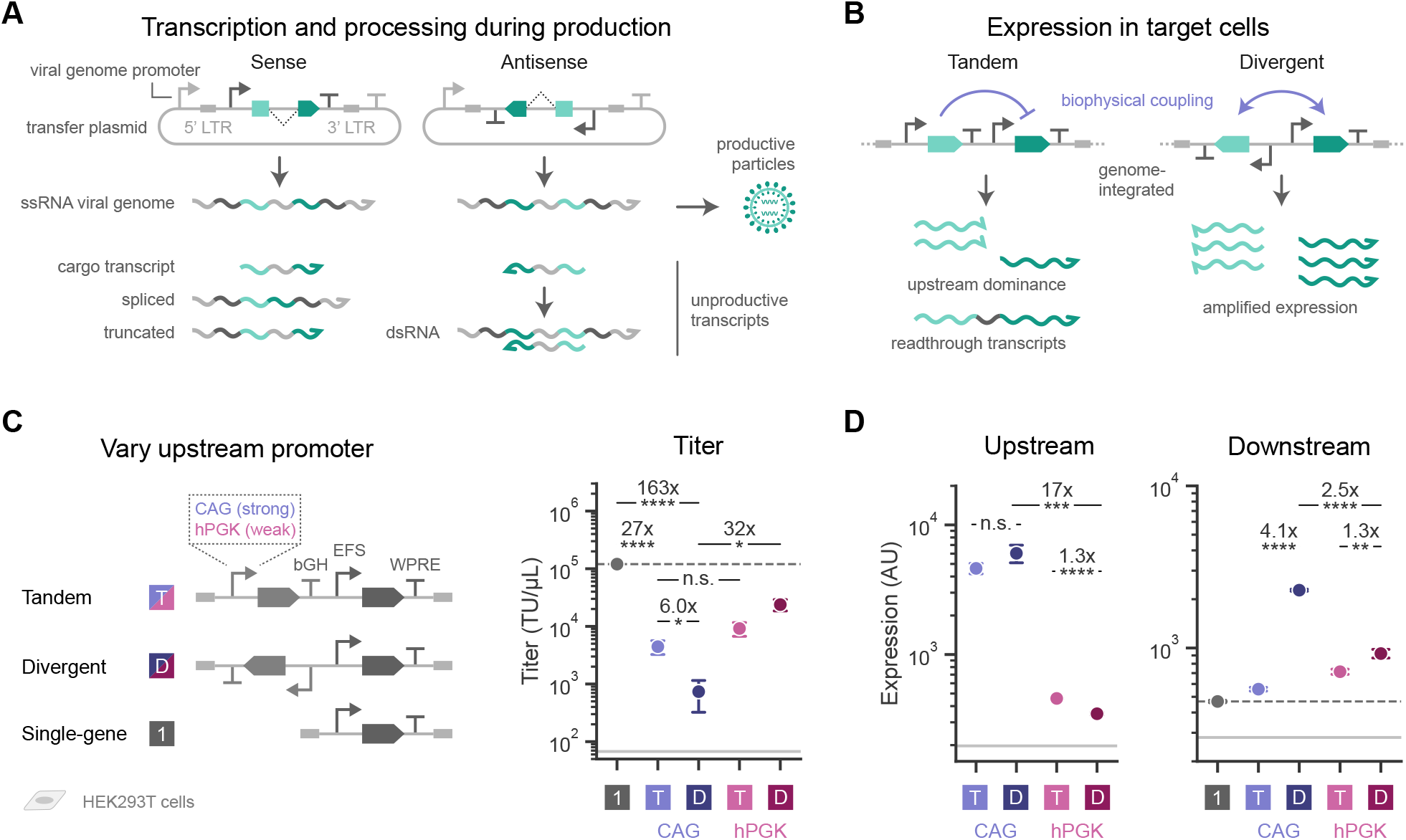
Promoter strength and gene syntax define tradeoffs in two-gene vectors. A. Schematic of the RNA species transcribed in producer cells for vectors with cargo on the sense or antisense strand relative to the viral genome. The transfer plasmid expresses the viral genome, as well as cargo transcripts and spliced or truncated forms of the viral transcript. The viral genome can be packaged into productive particles, while the other species are unproductive, meaning they do not form viral particles that integrate the intended sequence into target cells. Additionally, antisense cargo transcripts can form double-stranded RNA (dsRNA) by binding to the viral genome, potentially inhibiting proper packaging. B. Expression in target cells is influenced by biophysical coupling defined by the syntax of the encoded genes, resulting in attenuated or amplified expression. C. Schematic of two-gene vectors with tandem or divergent syntax that express a fluorescent protein from the strong, synthetic CAG promoter or from the relatively weak human phosphoglycerate kinase (hPGK) promoter, and a second fluorescent protein from a downstream EFS promoter. These vectors are compared to a single-gene vector containing only the downstream gene from Fig. **1**. The upstream gene includes the bovine growth hormone (bGH) polyadenylation signal (PAS), and the downstream gene includes the woodchuck hepatitis virus post-transcriptional regulatory element (WPRE) in the 3’ UTR. Viral titer is in transducing units (TU) per µL (log scale). D. Expression of the upstream and downstream genes for each vector in HEK293T cells. Expression is the geometric mean fluorescence in arbitrary units (AU, log scale). Solid light gray lines represent titers calculated for untransduced cells or expression gates. Dashed dark gray lines depict values for the single-gene vector for reference. Points represent means ±standard error for *n* ≥3 biological replicates. Statistics are two-sided Student’s t-tests, n.s. *p* ≥ 0.05, * *p <* 0.05, ** *p <* 0.01, *** *p <* 0.001, **** *p <* 0.0001. Annotations show the fold change between indicated points.

To build two-gene vectors, we added to the single-gene vectors an upstream gene expressing a fluorescent protein under control of the strong, synthetic CAG promoter. We encoded the two genes with tandem or divergent syntax and measured both titer and expression (Figs. **2C**, **S1A, S1B**). Compared to the best-producing single-gene vector, the two-gene tandem vector generates 27-fold lower titer with either the EFS or EF1α promoter in the downstream gene (Figs. **2C**, **S1B**). For divergent syntax, titer drops an additional six-fold, a decrease of more than two orders of magnitude compared to the best single-gene vector (Fig. **2C**). We hypothesize that with divergent syntax, strong antisense transcription from the upstream gene interferes with production of the viral genome.

In contrast, when transduced at constant MOI, the divergent vector results in four-fold higher mean expression of the downstream gene compared to the tandem vector and at least 1.8-fold higher expression than the single-gene vectors (Figs. **2D**, **S1B**), matching predictions of transcriptional amplification. For all designs, the EF1α promoter produces higher expression of the downstream gene compared to the EFS promoter, as expected (Fig. **S1B**). Finally, expression of the CAG-driven upstream gene is similar for the divergent and tandem vectors (Figs. **2D**, **S1B**). Thus, there exists a tradeoff between titer and expression level for these vectors, with the divergent design expressing most strongly but producing least efficiently.

We hypothesized that replacing CAG with a weaker upstream promoter could restore titer for the divergent design. Indeed, using a promoter derived from the human phosphoglycerate kinase gene (hPGK), which drives lower expression^48^, titers of divergent two-gene vectors increase by more than an order of magnitude (Figs. **2C**, **S1B**). Meanwhile, the titer of tandem designs do not increase, and none of the two-gene designs are produced as efficiently as the best single-gene vector. However, as expected, replacing CAG with hPGK reduces expression of the upstream gene for both syntaxes (Figs. **2D**, **S1B**), further underscoring the tradeoff between viral titer and upstream expression.

### An inducible upstream promoter resolves tradeoffs in vector performance

To overcome these tradeoffs, an ideal vector would provide strong expression in target cells while limiting interference in producer cells. An inducible promoter fulfills these criteria. In the absence of inducer, the upstream gene generates low, basal levels of transcription in producer cells, minimally disrupting production of the viral transcript (Fig. **3A**). In target cells, addition of the inducer turns on the gene of interest after transduction. We chose the widely used Tet-ON system for the inducible gene, in which the TRE3G promoter is activated by the small molecule doxycycline (dox). Binding of dox to the reverse tetracycline transactivator (rtTA) protein allows rtTA to bind to the TRE3G promoter, inducing transcription. To measure performance of these vectors, we separately integrated an rtTA-expressing cassette into HEK293T cells. For the divergent vector, replacing the hPGK promoter with TRE3G increases the viral titer three-fold, nearing the level of the single-gene vector and remaining higher than the titer for the tandem vector (Fig. **3B**). Expression of the upstream gene is an order of magnitude larger with TRE3G upon induction than with hPGK for both syntaxes, reaching the level achieved by the CAG promoter (Fig. **3B**). Across all metrics, the inducible divergent vector matches or exceeds performance of the corresponding single-gene and tandem vectors, resolving the tradeoff between titer and expression.

**Figure 3.**
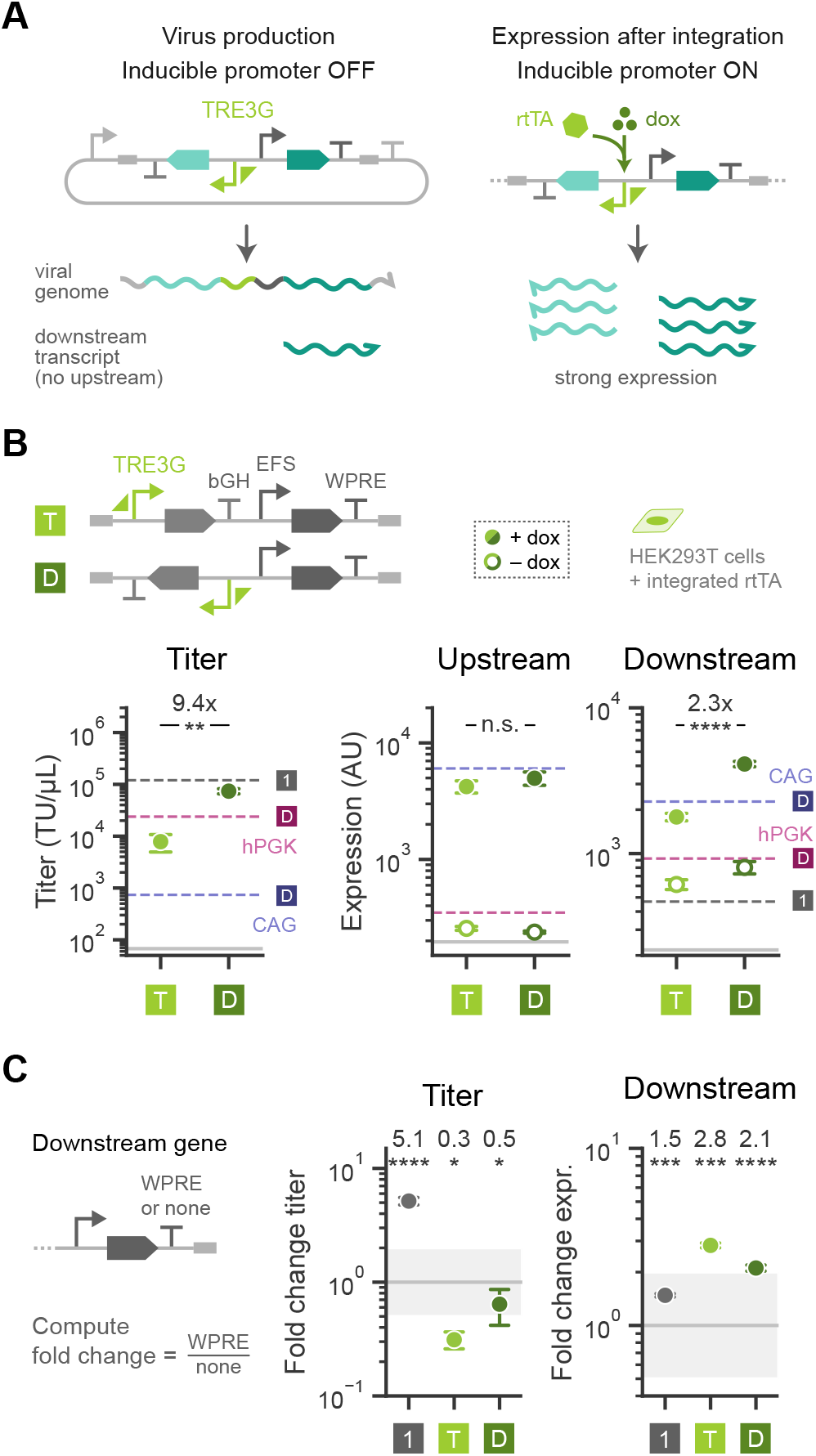
An inducible upstream promoter can resolve tradeoffs in vector performance. A. Schematic of transcripts produced from a divergent vector with the upstream, antisense gene driven by a Tet-ON inducible promoter (TRE3G). In producer cells cultured without inducer, the viral genome is transcribed from the transfer plasmid, and some unproductive downstream transcripts may also be produced. After integration in target cells, addition of the inducer doxycycline (dox) and the reverse tetracycline transactivator (rtTA) protein leads to strong expression of the inducible gene, along with strong expression of the downstream gene due to biophysical coupling. B. Two-gene vectors with an inducible upstream gene and tandem (T) or divergent (D) syntax were delivered to HEK293T cells containing a separately integrated rtTA cassette. Titer is in transducing units (TU) per µL (log scale), and expression is the geometric mean fluorescence in arbitrary units (AU, log scale). Cells were treated with 1 µg/mL dox (filled points) or untreated (open points). Solid light gray lines represent titers calculated for untransduced cells or expression gates. Dashed lines depict values for several vectors from Fig. **2** for reference. Annotations show the fold change between indicated points. C. Performance of vectors in B is compared to vectors lacking WPRE in the 3’ UTR of the downstream gene. Titer and expression of the downstream gene for the vectors with WPRE are normalized to values for vectors without WPRE, plotted on a log scale. Solid gray line indicates a fold change of 1, and gray shading spans a two-fold change in either direction (0.5 to 2) for reference. Annotations show the values of the points, and statistics compare unnormalized values. Points represent means ±standard error for *n*≥ 3 biological replicates. Statistics are two-sided Student’s t-tests, n.s. *p* ≥ 0.05, * *p <* 0.05, ** *p <* 0.01, *** *p <* 0.001, **** *p <* 0.0001.

Having identified a promoter combination that optimizes viral titer and expression of both cargoes, we next sought to understand how elements in the 3’ UTR such as a PAS affect vector performance. In our initial testing, we found that for tandem syntax, the choice of upstream PAS unexpectedly has little impact on viral titer and upstream expression (Fig. **S2A**). However, for the divergent vectors, including the bGH PAS moderately improves both metrics, though the presence of the bGH PAS on both the upstream and downstream genes abolishes virus production (Fig. **S2A**). For the downstream gene, our initial vector designs contained the woodchuck hepatitis virus post-transcriptional regulatory element (WPRE) in the 3’ UTR without a subsequent PAS. WPRE is commonly included in viral vectors to enhance expression, supporting several RNA processing steps including nuclear export and transcript stability^50,51^. Consistent with these findings, addition of the WPRE results in about a two-fold increase in expression of the downstream gene for both single-gene and inducible two-gene vectors (Figs. **3C**, **S2A, S2B**). Across all vectors, WPRE in combination with a subsequent PAS has mixed effects on titer (Figs. **3C**, **S2A, S2B**). In particular, the wide variation in titer for vectors with the EF1α promoter may suggest the appearance of emergent effects between 3’ elements and the EF1α intron. Overall, WPRE consistently affords a small boost in expression of the downstream gene with small effects on titer, and addition of a subsequent PAS does not broadly improve vector performance.

### “All-in-one” vectors with divergent syntax offer high titer and inducible control

While an inducible upstream gene elicits both high titer and strong expression, this design relies on the presence of the associated activator protein. Instead of co-transducing a separate vector containing the activator, which may reduce overall delivery efficiency, the activator can be encoded within the constitutively expressed gene on the same vector to generate an “all-in-one” design^52^. We constructed “all-in-one” Tet-ON vectors with tandem and divergent syntaxes, expressing rtTA alongside the downstream fluorescent protein separated by a “self-cleaving” 2A peptide (Fig. **4**). As expected, both titer and expression of each gene match values for the inducible two-gene vectors lacking the rtTA (Fig. **S3**). In the absence of inducer, expression of the upstream gene is minimal (Fig. **S3B**). Both syntaxes support robust induction (Figs. **4**, **S3B**), consistent with our prior investigation of gene syntax in diverse cell types^40^. However, with these genetic parts, the divergent design increases titer five-fold over the tandem vector (Figs. **4**, **S3A**).

**Figure 4.**
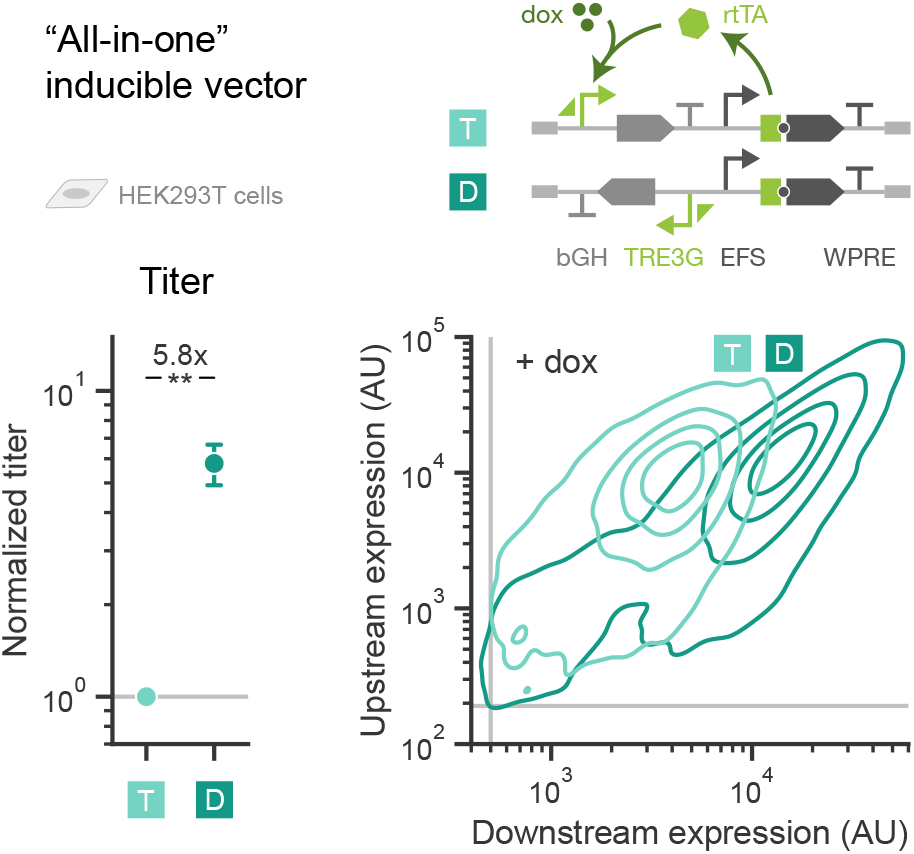
“All-in-one” vectors with divergent syntax offer high titer and inducible control. An “all-in-one” inducible vector contains both an inducible gene and a constitutive gene that expresses the required activator, which can be arranged with tandem (T) or divergent (D) syntax. Here, the dox-inducible TRE3G promoter (Tet-ON system) controls expression of a fluorescent protein in the upstream gene, and in the downstream position EFS drives expression of rtTA and a second fluorescent protein, separated by a “self-cleaving” 2A peptide. The upstream gene includes the bGH PAS, and the downstream gene includes WPRE. **Left:** Viral titer is shown normalized to the tandem vector for each batch of virus. Points represent means ±standard error for *n* ≥3 biological replicates. Statistic is a two-sided Student’s t-test, ** 0.05 ≤*p <* 0.01. **Right:** 2D density distributions depict upstream and downstream expression via fluorescence in arbitrary units (AU, log scale) for a representative biological replicate of HEK293T cells transduced with each vector and treated with 1 µg/mL dox. Populations are gated on cells expressing both genes. Solid gray lines show expression gates.

To explore the generality of the “all-in-one” design, we varied the upstream PAS and downstream promoter. Divergent vectors have higher upstream expression but similar titer with a bGH PAS versus no PAS, while tandem designs have higher titer but similar expression with no PAS for downstream EFS and hPGK promoters (Fig. **S3**). With a downstream EF1α promoter, trends are less clear for the tandem vectors, similar to the variation observed for the single-gene vectors (Fig. **S2A**). Additionally, for both divergent and tandem vectors, expression of the downstream gene is similar with either the EFS or EF1α promoter (Fig. **S3B**). This stands in contrast to the previous and expected observation of higher expression with EF1α in single-gene and two-gene vectors lacking the rtTA (Figs. **1C**, **S1B, S2A, S2B**). These results suggest that the effects of the EF1α promoter may be less predictable than those of promoters lacking an intron (e.g., EFS, hPGK). Nevertheless, the divergent vector with the bGH PAS in the upstream gene is the best-performing design for each downstream promoter.

### The divergent “all-in-one” design effectively encodes diverse RNA-based gene circuits

Although some tandem “all-in-one” designs perform well, we hypothesized that inclusion of RNA devices such as microRNAs, ribozymes, and splicing switches in the sense direction of the viral transcript could alter RNA processing and decrease production efficiency. In contrast, encoding these elements on the antisense strand (i.e., in the upstream gene with divergent syntax) would preclude device activity during production. To test this hypothesis, we first delivered ComMAND, a post-transcriptional circuit that controls for gene dosage^53^. ComMAND uses a synthetic microRNA within an intron of the output coding sequence to target a matched microRNA target site in the 3’ UTR of the same transcript, resulting in an incoherent feedforward loop that buffers variation in expression. We expressed the circuit in the upstream gene of the best-performing divergent and tandem “all-in-one” vectors and compared these designs to vectors containing only the output coding sequence (denoted as “base gene”) and to an open-loop circuit containing the microRNA and an orthogonal target site (Figs. **5A**, **S4**). Compared to the base gene vector, both the open-loop circuit and ComMAND decrease titer for vectors with tandem syntax (Figs. **5A**, **S4A**). In contrast, for divergent syntax, titer remains the same across circuits, resulting in a 50-fold greater titer for the ComMAND vector with divergent syntax over the tandem design. Meanwhile, syntax minimally impacts circuit function: both the tandem and divergent designs show comparable reduction in expression variability with ComMAND relative to the base gene and open-loop circuit (Figs. **5A**, **S4B**–**S4D**).

**Figure 5.**
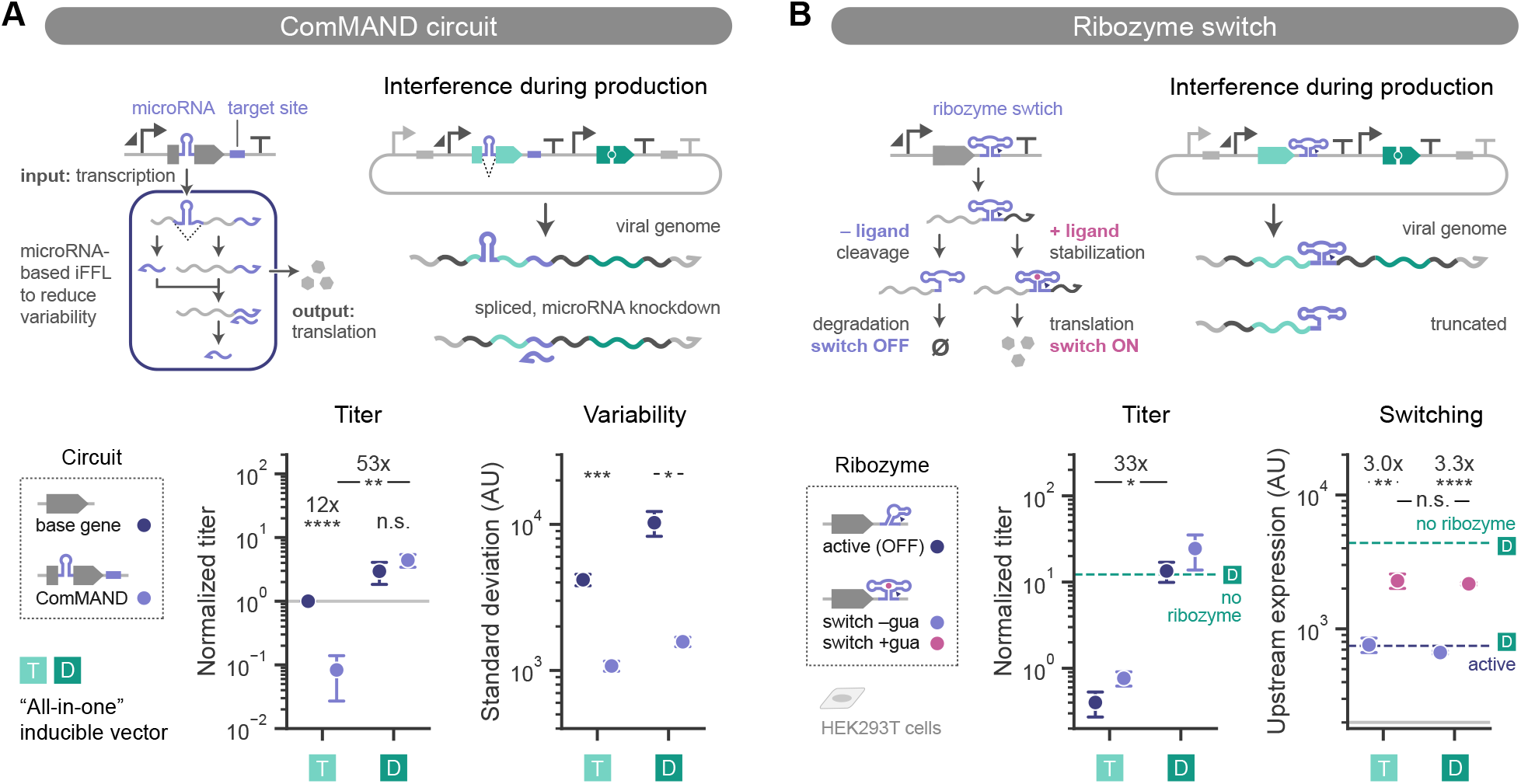
The divergent “all-in-one” inducible vector effectively delivers RNA-based circuits. A. ComMAND is a compact, post-transcriptional circuit that reduces dosage-associated variability in expression. ComMAND uses a synthetic microRNA within an intron of the output coding sequence to target a matched microRNA target site in the 3’ UTR of the same transcript, resulting in an incoherent feedforward loop. Because ComMAND operates at the RNA level, circuit activity can interfere with virus production when encoded on the sense strand of the viral genome. “All-in-one” inducible vectors as in Fig. **4** were constructed with ComMAND (light purple) or the output gene alone (base gene, dark blue) as the upstream cargo. **Left:** Titer is normalized to the base gene vector with tandem syntax for each batch of virus (log scale). **Right:** Standard deviation of circuit (upstream gene) expression measured via fluorescence in arbitrary units (AU, log scale). B. A ribozyme switch, consisting of a ligand-responsive RNA aptamer fused with a self-cleaving ribozyme, controls expression of an output gene when placed in the 3’ UTR. In the absence of ligand, the ribozyme switch cleaves, leading to degradation of the mRNA and a reduction in protein levels (OFF). Binding of the ligand changes the folding of the switch to minimize ribozyme cleavage and restore protein expression (ON). When encoded on the sense strand of the viral genome, ribozymes can interfere with virus production by cleaving and truncating transcripts. “All-in-one” inducible vectors as in Fig. **4** were constructed with a guanine (gua)- responsive ribozyme switch or a constitutively active ribozyme added to the upstream gene. **Left:** Titer is normalized to a vector lacking a ribozyme (not shown) for each batch of virus (log scale). **Right:** Upstream expression is the geometric mean fluorescence (AU, log scale) of cells transduced with vectors containing the ribozyme switch and treated without (light purple) or with (pink) 100 µM guanine. Solid light gray line shows the expression gate. Dashed teal lines depict values for a divergent vector lacking a ribozyme, and dashed dark blue line represents the value for the divergent vector with the constitutively active ribozyme. To compare optimal designs for each syntax, tandem vectors lack a PAS on the upstream gene, while divergent vectors include a bGH PAS. In all plots, HEK293T cells were transduced and treated with 1 µg/mL dox. Points represent means ±standard error for *n* ≥3 biological replicates. Statistics are two-sided Student’s t-tests, n.s. *p* ≥0.05, * *p <* 0.05, ** *p <* 0.01, *** *p <* 0.001, **** *p <* 0.0001. Annotations show the fold change between indicated points.

To assess the generality of this syntax-specific performance, we constructed tandem and divergent “all-in-one” vectors expressing a ligand-responsive ribozyme switch^54^ in the 3’ UTR of the upstream gene (Figs. **5B**, **S5**). In the absence of ligand, the ribozyme element in the switch self-cleaves, decreasing transcript stability and thus protein expression. Binding of the ligand to an aptamer in the switch alters the conformation of the ribozyme, reducing cleavage rates and increasing protein expression. As controls, we replaced the ribozyme switch with a constitutively active ribozyme (protein expression OFF) or a mutant, minimally cleaving ribozyme (protein expression ON). Analogous to our observations for ComMAND, titer remains similar across vectors with divergent syntax but decreases with increasing ribozyme activity for tandem syntax (Figs. **5B**, **S5A**). Furthermore, upstream expression and switch performance are the same between syntaxes (Figs. **5B**, **S5C**). Together, divergent syntax improves production of lentiviral vectors containing RNA devices such as microRNAs and ribozymes while maintaining function of the RNA-based circuits in target cells.

### Divergent syntax supports small-molecule tuning of expression from a splicing switch

Observing that both syntaxes lead to strong circuit performance for ComMAND and the ribozyme switch, we next explored how the performance of a splicing switch is affected by lentiviral delivery. Encoded between exons of a gene of interest, the splicing switch consists of introns flanking an alternative exon with a premature stop codon. In the absence of the ligand guanine, splicing includes the alternative exon in the mature mRNA, leading to truncated protein (expression OFF). Binding of guanine to an aptamer in one of the introns alters the RNA structure and changes the splicing pattern to exclude the alternative exon, resulting in production of full-length protein (expression ON) (Fig. **6A**). The amount of guanine sets the ratio of these splice products in cells to tune protein levels. We found that “all-in-one” vectors encoding this RNA device have similar titers for the best tandem and divergent designs, though addition of an upstream bGH PAS with tandem syntax abolishes virus production (Figs. **6A**, **S6A**). However, the tandem vector without an upstream PAS has no switch function, despite its efficient delivery as evidenced by expression of the downstream gene (Fig. **6A**, Figs. **S6A**–**S6C**). Target cells transduced with this vector express the upstream gene in the absence of guanine, and addition of guanine does not change these levels. In contrast, the divergent vector results in upstream expression that is minimal in the absence of guanine and that increases 100-fold upon induction (Figs. **6A**, **S6A, S6C**), as intended for the switch.

**Figure 6.**
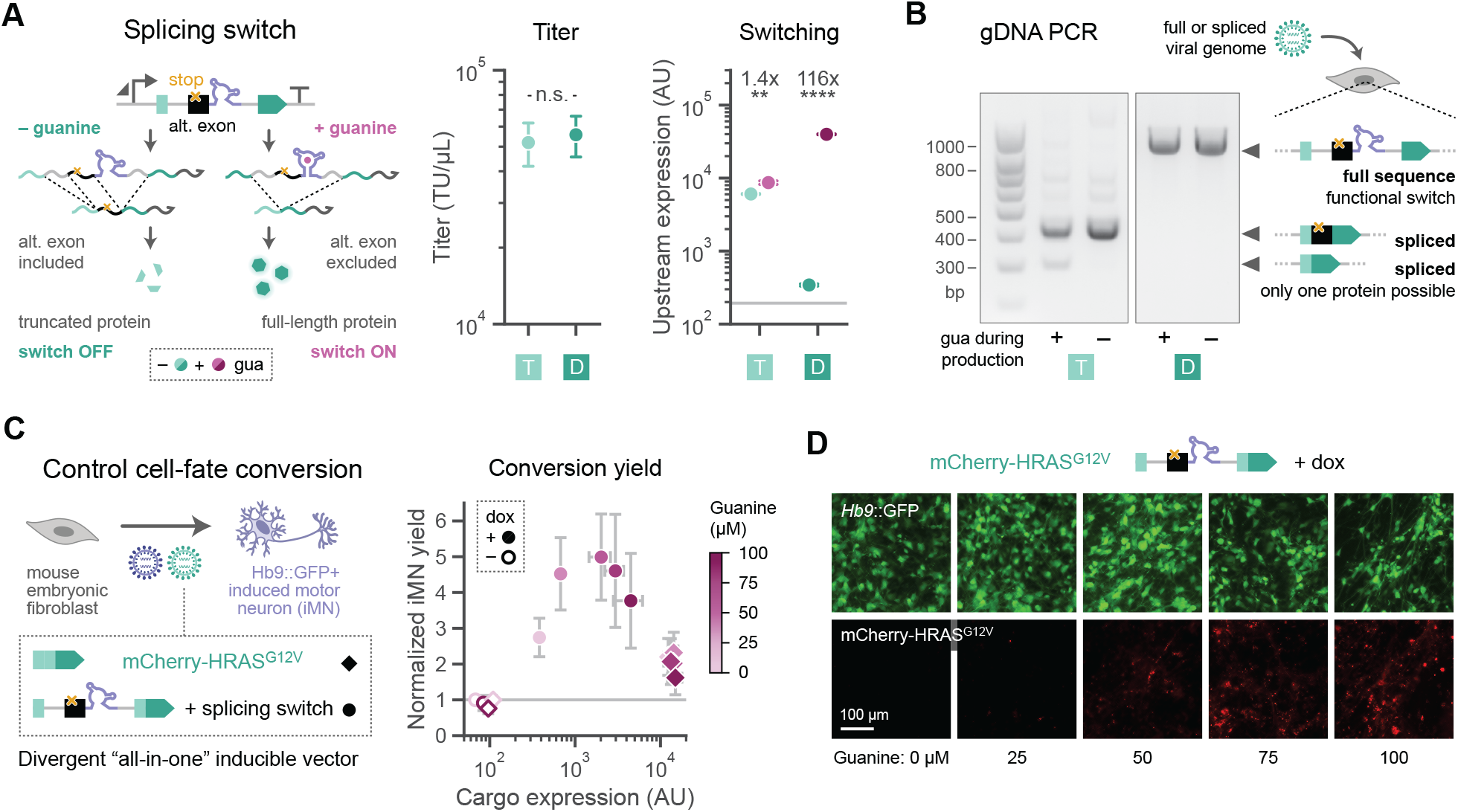
Divergent syntax supports small-molecule tuning of expression from a splicing switch. A. A guanine-responsive splicing switch consists of two introns and an alternative exon (alt. exon) containing a stop codon. Addition of the splicing switch within the coding sequence of a gene of interest enables tunable control of protein expression. In the absence of guanine (gua), the main spliced isoform includes the alternative exon, leading to truncated protein product. In the presence of guanine, the RNA aptamer changes structure and alters the splicing pattern to exclude the alternative exon and enable production of full-length protein. Switches are encoded within the upstream gene of the best “all-in-one” inducible vectors identified previously (Fig. **4**). The tandem vector (T) does not include a PAS on the upstream gene while the divergent vector (D) includes the bGH PAS. **Left:** Titer is measured in transducing units (TU) per µL (log scale). **Right:** Upstream expression is geometric mean fluorescence in arbitrary units (AU, log scale) for HEK293T cells treated with 1 µg/mL dox and without (teal) or with (pink) 100 µM guanine. Solid light gray line shows the expression gate. B. Gel depicts PCR products amplified from genomic DNA (gDNA) of cells transduced with vectors containing the splicing switch. Producer cells were cultured with or without 100 µM guanine during virus production. The amplicon spans the site where the splicing switch was added. Lengths of the ladder bands are labeled in base pairs (bp), and the most prominent bands in the conditions are annotated with their predicted splice products. The uncropped gel is displayed in Fig. **S6E**. C. Schematic of direct conversion of primary mouse embryonic fibroblasts to *Hb9*::GFP+ induced motor neurons (iMNs) via viral delivery of conversion factors. *Hb9::GFP* is a transgenic reporter of iMN fate used to identify converted cells. The splicing switch regulates expression of the oncogenic HRAS mutant HRAS^G12V^, which is fused to the fluorescent protein mCherry and delivered via the the best-performing divergent “all-in-one” inducible vector. Vectors contain the splicing switch within the mCherry coding sequence (circles) or lack the switch (diamonds) and were transduced into mouse embryonic fibroblasts alongside the remaining conversion factors. Cells were treated with 1 µg/mL dox (filled points) or without (open points), and additionally treated with 0–100 µM guanine for the duration of the conversion process. At 14 days post-transduction, iMN yield was quantified as the number of cells expressing *Hb9::GFP* per cell seeded, normalized to the same count for the untreated condition (neither dox nor guanine added) for each vector for each biological replicate. Cargo expression is the geometric mean of mCherry fluorescence for the iMN population (AU, log scale). D. Images show fluorescence of *Hb9*::GFP (top) and mCherry-HRAS^G12V^ (bottom) at 14 days post-transduction for cells transduced with the vector containing the splicing switch, treated with dox, and treated with varying concentrations of guanine. Scale bar represents 100 µm. Points represent means ±standard error for *n*≥ 3 biological replicates. Statistics are two-sided Student’s t-tests, n.s. *p*≥ 0.05, * *p <* 0.05, ** *p <* 0.01, *** *p <* 0.001, **** *p <* 0.0001. Annotations show the fold change between indicated points.

We hypothesized that the lack of inducibility for the tandem design results from integration of an incomplete splicing switch (Fig. **S6D**). To investigate the integrated sequence, we used PCR to amplify genomic DNA isolated from cells transduced with each vector. The single amplicon for the divergent vector matches the expected size for the construct containing the full-length splicing switch, while conditions with the tandem vector have multiple amplicons (Figs. **6B**, **S6E**). For the tandem vector, the amplicon at the full-length size is absent, and the most prominent band on the gel matches the splice product containing the alternative exon but lacking the intronic regions. This suggests that the viral transcript undergoes splicing in producer cells, packaging sequences that lack essential switch components. To test this hypothesis, we treated producer cells with guanine during virus production, reasoning that this would activate the splicing switch and generate viral transcripts without the alternative exon (Fig. **S6D**). Indeed, performing the same PCR on gDNA from cells transduced with these vectors resulted in an amplicon corresponding to the protein coding sequence without the entire switch (Figs. **6B**, **S6E**). Further, for these conditions, expression increases in target cells without restoring responsiveness to guanine (Figs. **S6A**–**S6C**). Thus, during production of tandem vectors, the splicing switch is active and changes the splice products of the viral transcripts that are packaged into particles. Conversely, addition of guanine during production of the divergent vector has no impact on gDNA PCR amplicons, expression, or switch activity (Figs. **6A, 6B**, **S6**). Thus, divergent syntax supports production and delivery of a functional splicing switch on a lentiviral vector.

Having identified a high-titer, guanine-responsive vector design, we used the splicing switch to titrate expression of a conversion factor during direct conversion of primary mouse embryonic fibroblasts to induced motor neurons (iMNs) (Figs. **6C**, **S7A, S7B**). High rates of conversion rely on delivery of a high-efficiency cocktail that combines with neuronal transcription factors to drive cells to adopt a motor neuron identity^55–57^. Within this cocktail, we had previously identified that an intermediate level of the HRAS mutant HRAS^G12V^ is optimal for conversion^58^. To titrate expression, we had used an array of techniques including varying lentiviral dosage, cassette design, promoter usage, and promoter setpoint editing^58,59^. In contrast, we hypothesized that the splicing switch would allow us to easily modulate HRAS^G12V^ expression via guanine addition while maintaining identical transduction conditions and a constant MOI. As a control, we also constructed a vector expressing HRAS^G12V^ without the splicing switch. As desired, titrating the amount of guanine unimodally shifts expression of the cargo over an order of magnitude in converting cells (Figs. **6C, 6D**, **S7C**–**S7E**). For the same conditions, we measured conversion yield by counting the number of cells expressing *Hb9::GFP* per cell seeded and normalizing to controls without overexpression of HRAS^G12V^. Matching previous findings^58^, we determined that intermediate concentrations of guanine and levels of HRAS^G12V^ expression optimize iMN yield (Figs. **6C, 6D**, **S7E**). Together, the divergent “all-in-one” vector incorporating the splicing switch provides a robust and practical approach for tuning expression of a functional cargo in primary cells.

## Discussion

In this work, we sought to define design rules for constructing lentiviral vectors to enhance delivery and control of genetic cargoes in human cells. We focused on vectors containing two transcriptional units that incorporate RNA devices and circuits, as these tools offer sophisticated control over cargo expression. Measuring functional titer reveals effects of vector design across the virus lifecycle, from production and packaging of the viral transcript to integration in target cells. Production efficiency is crucial for downstream applications of lentiviral vectors, as it impacts manufacturing scale-up. We observed tradeoffs between viral titer and functional expression for some designs. We addressed this challenge by expressing one of the genes from an inducible promoter on the antisense strand of the viral transcript (i.e., with divergent syntax), and constructed effective “all-in-one” vectors that also express the required transactivator. These vectors are similar to some reported in the literature^60^. However, previous studies do not compare gene syntax in combination with genetic parts. Studies examining the affects of varying genetic parts primarily focus on expression in target cells and often lack characterization of production efficiency. We aimed to systematically fill this knowledge gap to facilitate predictable vector design.

In the best-performing vectors, we encoded a microRNA-based circuit, ribozyme switch, and splicing switch, finding that the “all-in-one” design with divergent syntax affords both high titer and functionality for these cargoes, while vectors with tandem syntax do not. Our results suggest that when encoded on the sense strand, these RNA devices primarily reduce titer by preventing formation of particles with intact viral transcripts. Similarly, these designs may compromise functionality by altering the sequence of packaged transcripts. As lentiviral vectors are increasingly used to deliver gene therapies *in vivo*, integration of aberrant sequences presents clinical risks. In contrast, when encoded on the antisense strand with divergent syntax, we find that these RNA devices minimally impact virus production, express well in target cells, and retain their characteristic functions. We anticipate that our “all-in-one” vectors offer an effective framework for delivering other sequences that may interfere with processing of the viral transcript.

We expect our best-performing vector design to generalize beyond the cargo, genetic parts, and cell types tested here. As a demonstration, our best-performing design expresses a functional cargo, a dose-dependent cell-fate conversion factor, suggesting that the vector can deliver other therapeutic payloads. Nevertheless, further testing across diverse primary human cell types is required for downstream applications. Encoding native genes in the vector may affect titer by introduction of sequences that affect RNA processing, such as cryptic splice sites^31–35^. Comparison to well-defined lentiviral vectors may reveal hidden features within cargo sequences and offer a design alternatives. Further, cargo size may also influence vector performance, warranting future work. As we expand our designs to other sequences, we envision replacing the bacterial-derived Tet-ON system with humanized components^61,62^ and FDA-approved inducers, such as the synZiFTR platform^63^, for greater clinical relevance. We may also consider exchanging other elements for human-derived sequences. Though viral in origin, the WPRE sequence is often included in lentiviral vectors, as it reliably boosts expression^50,51^ with minimal effect on titer, which our work supports, and mitigates transgene silencing^47^. Although we did not test vectors encoding this element in the 3’ UTR of the upstream gene, we hypothesize that two copies of WPRE in divergent syntax may abolish vector production, as observed for two bGH PAS sequences. Finally, while inclusion of an upstream PAS consistently increases titer and expression for the divergent vectors we tested, tandem vectors performed as well or better without an upstream PAS, though it is possible that other PAS elements may behave differently.

Fundamentally, our approach leverages an inducible promoter that does not express in producer cells to maintain high titer, and that elicits strong expression in target cells upon induction. Beyond the Tet-ON system used here, we expect that cell state-responsive promoters that are transcriptionally active in target cells but not producer cells would fulfill these criteria^64–67^. Using promoters that recruit endogenous transcriptional machinery would eliminate the need for an exogenous transactivator, reducing cargo size and avoiding potential side effects of synthetic systems. One potential challenge to this approach is that cell state-responsive promoters may have more “leaky” expression outside of the intended cell state, which may be exacerbated by nearby transcription in divergent syntax^37^. We did not observe expression in uninduced conditions for genes regulated by the tightly controlled Tet-ON system. However, even with tandem syntax, we observed an unexpected increase in expression of the downstream EFS-driven gene upon induction of the adjacent TRE3G promoter, though not to the same degree as with divergent syntax. Potentially, the high local concentration of transcriptional machinery recruited to the inducible gene enhances expression of the neighboring gene regardless of syntax. This effect may only be apparent for very strong inducible promoters like TRE3G used here^48^, or it may be specific to EF1α-derived promoters, as vectors with hPGK did not exhibit this behavior. Nevertheless, effects of the inducible gene on the neighboring constitutive gene could disrupt expression of functional cargoes with narrow expression windows. As yet, the easiest way to robustly “insulate” expression of one gene from another is to deliver them on separate vectors, at the cost of delivery efficiency. Future studies could determine the generality of this coupling and devise new insulation approaches.

Together, this work illuminates key design principles for constructing two-gene lentiviral vectors encoding RNA-based circuits. Our results highlight the pitfalls of common designs and offer strategies for robust manufacturability and functionality. Optimizing for both of these properties will ensure that vectors are efficacious and support scalable production, advancing toward feasible clinical applications. Delivering compact, tunable, and modular RNA-based circuits on lentiviral vectors will unlock avenues for precise control in gene and cell therapies.

## Materials and methods

### Cloning

All lentiviral transfer plasmids were constructed using a multi-level Golden Gate cloning scheme described in Peterman *et al*.^48^, with the third-generation Lenti-X1 lentiviral backbone derived from Addgene #17297. ComMAND circuit^53^ plasmids include Addgene #235268, #235319, and #235320, and additional plasmids were derived from these. The plasmids containing the constitutively active and mutant ribozymes^10^ were derived from Addgene #131744 and #131745, and the ribozyme switch was ordered as a gBlock from Azenta/Genewiz using the sequence from Mustafina *et al*.^54^. The splicing switch was a gift from Shiva Razavi, Jonathan C. Chen, and James J. Collins at MIT. Plasmids containing the splicing switch were derived from this vector. The plasmids containing HRAS^G12V^ were derived from Addgene #246340 (ref.^59^). Sources for other genetic parts can be found in Peterman *et al*.^48^. Key plasmids will be deposited at Addgene, and other plasmids will be made available upon request.

## Cell culture

### Cell lines

HEK293T cells (ATCC, CRL-3216), Lenti-X HEK293T cells (Takara Bio, 632180), and Plat-E retroviral packaging HEK293T cells (Cell Biolabs, RV-101) were cultured using DMEM (Genesee Scientific, 25-500) plus 10% FBS (Genesee Scientific, 25-514H) and incubated at 37°C with 5% CO_2_. For routine passaging, seeding, and preparation for flow cytometry, HEK293T cells were dissociated using 0.25% Trypsin-EDTA (Genesee Scientific, 25-510) diluted in PBS (Sigma-Aldrich, P4417-100TAB) and incubated at 37°C for four minutes, then quenched with an equal volume of DMEM + 10% FBS. Cells were passaged about every three days to maintain consistent growth before experiments. Plat-E cells were selected using 10 µg/mL blasticidin and 1 µg/mL puromycin every three passages. Cells were tested for mycoplasma.

### Primary mouse embryonic fibroblasts

Primary mouse embryonic fibroblasts were isolated as described in Wang *et al*.^56^. C57BL/6 mice (The Jackson Laboratory, 000664) were mated with B6CBAF1 mice bearing the transgenic *Hb9::EGFP* reporter (The Jackson Laboratory, 005029), and embryos were harvested at E12.5-E14.5 under a dissection scope. Embryo heads and internal organs were removed, then razors were used to break up the tissue with the addition of 0.25% Trypsin-EDTA. One or two embryos were processed simultaneously. After five minutes, the solution was quenched with DMEM + 10% FBS, spun down, and resuspended in fresh Trypsin-EDTA. After triturating, the solution was again quenched DMEM + 10% FBS and spun down. The resulting cells were resuspended in DMEM + 10% FBS and passed through a 40-µm filter, then plated on 10-cm dishes coated with 0.1% gelatin (Sigma-Aldrich, G1890-100G), one dish per embryo (passage 0). Cells were incubated at 37°C with 5% CO_2_. Once cells reached ∼80% confluence (after two to three days), they were dissociated using 0.25% Trypsin-EDTA and passaged 1:3 onto fresh, gelatin-coated 10-cm dishes (passage 1). During passaging, a subset of cells were removed, expanded, and tested for mycoplasma. Once confluent (after two to four days), cells were dissociated using 0.25% Trypsin-EDTA, cryopreserved in 10% DMSO + 90% FBS, and stored in liquid nitrogen. For experiments, vials of cryopreserved passage 1 cells were thawed into DMEM + 10% FBS in T75 flasks coated with 0.1% gelatin and allowed to recover for one to two days before seeding.

### Lentiviral vector production

Lenti-X HEK293T cells were seeded at 7.5 million cells per well in 10-cm dishes coated with 0.1% gelatin. The following day (day one), the transfer plasmid, packaging plasmid (psPAX2, Addgene #12260), and envelope plasmid (pMD2.G, Addgene #12259) were co-transfected using PEI (linear polyethylenimine, Fisher Scientific, AA4389603). Transfection mixes were prepared using a ratio of 4 µg PEI to 1 µg DNA. First, a master mix of PEI and KnockOut™ DMEM (Fisher Scientific, 10-829-018) was prepared and incubated for a minimum of ten minutes. This mixture was then added to prepared DNA mixes containing 6 µg of the transfer plasmid, 6 µg of the packaging plasmid, and 12 µg of the envelope plasmid per dish. These condition mixes were incubated for an additional 10 to 15 minutes and then added dropwise on top of the growth media in the 10-cm dishes. After six hours, the media was replaced with 6.5 mL of DMEM + 10% FBS with 25 mM HEPES (Sigma-Aldrich, H3375). On the following day (day two), the media was collected, stored at 4°C, and replaced with fresh 6.5 mL HEPES-buffered DMEM + 10% FBS. On day three, the media was again collected. The collected media was filtered through a 0.45-µm polyethersulfone (PES) filter (Thermo Scientific, CH4525-PES). To concentrate the virus, Lenti-X Concentrator (Takara Bio, 631232) was added to the filtered virus-containing media in a volume ratio of 3 parts media to 1 part concentrator, mixed gently, and left overnight at 4°C. On day four, the media was centrifuged at 1500x*g* at 4°C for 45 minutes. The supernatant was aspirated, and the resulting pellet was resuspended in 200 µL of DMEM + 10% FBS. Virus was stored at –80°C before use.

### Lentiviral vector transduction

On the day of transduction, HEK293T cells were dissociated and diluted to a concentration of 20,000 cells per well (50 µL/well) in DMEM + 10% FBS. Cells were combined with 5 µg/mL polybrene (hexadimethrine bromide, Sigma-Aldrich, H9268-5G) and the specified volume of virus (see below). Additional DMEM + 10% FBS was added to reach a total volume of 100 µL per well. The resulting cell, polybrene, and virus mixture was plated onto 96-well plates coated with 0.1% gelatin. The following day, the media was replaced with fresh DMEM + 10% FBS, with or without inducers according to the condition. Cells were prepared for flow cytometry two days later (three days post-transduction) by dissociating with Trypsin-EDTA. After centrifuging the dissociated cells at 500x*g* for five minutes, the cells were resuspended in PBS and transferred to a v-bottom plate for flow cytometry.

For transduction experiments to calculate functional titer, a two-fold serial dilution of concentrated virus was used, with six conditions per vector and 4 µL of concentrated virus per well in the highest condition. For transduction experiments using a constant multiplicity of infection (MOI), the volume of virus to use was calculated for each vector for each batch of virus from the titer data to achieve a MOI of 0.3. Namely, virus volume in µL, *v*, is a function of the number of cells being transduced, *c*, the MOI in transducing units (TU) per cell, *λ*, and the viral titer in TU/µL, *T* :

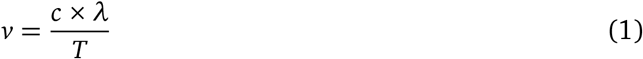

For vectors containing a Tet-ON inducible gene, 1 µg/mL dox (doxycycline, Sigma-Aldrich, D3447-500MG) dissolved in water was added to the media at one day post-transduction both for experiments to calculate titer and for experiments with a constant MOI. Experiments with a constant MOI also included conditions without dox. For vectors containing the ribozyme switch or the splicing switch, experiments with a constant MOI also included conditions treated with 100 µM guanine (from stocks of 25 mM dissolved in 0.2 N NaOH), with or without dox. Conditions without guanine were also included, using an equal volume of 0.2 N NaOH as a vehicle control instead.

### Direct conversion of primary mouse embryonic fibroblasts to induced motor neurons

#### Retroviral vector production

Conversion factors other than HRAS^G12V^ were delivered via retroviral vectors. These include LNI (Addgene #233154), a polycistronic cassette of Lhx3, Ngn2, and Isl1 neuronal transcription factors separated by 2A “self-cleaving” peptides, and SNAP-p53DD (Addgene #244168), a fusion and of the SNAPtag and an oncogenic p53 mutant lacking the DNA binding domain. To produce the retrovirus, Plat-E cells were seeded at 850,000 cells per well in 6-well plates coated with 0.1% gelatin. The next day, cells were transfected with 1.8 µg of transfer plasmid per well using PEI, then 24 hours later the media was replaced with 1.25 mL fresh HEPES-buffered DMEM + 10% FBS. On the following day, viral supernatant was collected and filtered through a 0.45-µm PES filter, replacing the media with 1.25 mL fresh HEPES-buffered DMEM + 10% FBS. Filtered viral supernatant was then immediately used to transduce mouse embryonic fibroblasts (described below). Viral supernatant collection, filtration, and transduction was repeated for a second day.

#### Transduction of primary mouse embryonic fibroblasts

Primary mouse embryonic fibroblasts were seeded at 5,000 cells per well in 96-well plates coated with 0.1% gelatin. The next day, media was replaced with fresh DMEM + 10% FBS plus 11 µL of each retrovirus (LNI and SNAP-p53DD) from the first collection. Cells were spinfected by centrifuging at 1500x*g* for 30 minutes. On the following day (defined as 0 days post-transduction), media was again replaced with fresh DMEM + 10% FBS plus 11 µL of each retrovirus from the second collection, with addition of 1 µL of concentrated HRAS^G12V^ lentivirus. Cells were again spinfected. After 24 hours, media was replaced with fresh DMEM + 10% FBS containing 1 µg/mL dox, 0–100 µM guanine, and/or vehicle controls, according to the condition. Conditions with inducer included these for the remainder of the experiment. At three days post-transduction, cells were switched into N3 media: DMEM/F-12 (Fisher Scientific, 21-331-020) supplemented with N-2 (Thermo Fisher Scientific, 17-502-048), B-27 (Thermo Scientific, 17-504-044), 1% GlutaMAX (Thermo Fisher Scientific, 35-050-061), 7.5 µM RepSox (a TGF-β inhibitor, Selleck Chemicals, S7223), and 10 ng/mL each of recombinant human BDNF (R&D Systems, 248-BDB-050), GDNF (R&D Systems, 212-GD-050), CNTF (R&D Systems, 257-NT-050), and FGF (Peprotech, 100-18B). Media was replaced with fresh N3 media every 2 to 3 days. At 14 days post-transduction, cells were imaged then dissociated for flow cytometry by incubating with DMEM/F12 plus 17 U/mL DNase (Worthington Biochemical, LK003172) and 167 U/mL papain (Worthington Biochemical, LK003178) at 37°C for 15 minutes. The cells were then centrifuged at 400x*g* for four minutes, resuspended in PBS, and transferred to a v-bottom plate for flow cytometry.

#### Amplification of genomic DNA from transduced cells

HEK293T cells were transduced with 4 µL of concentrated lentivirus per well in a 96-well plate as for transductions to calculate functional titer. At three days post-transduction, cells were dissociated using Trypsin-EDTA, spun down at 500x*g* for 5 minutes, and combined with 50 µL Cell Lysis Buffer (Cell Signaling Technology, 9803S) and 1 µL Proteinase K (New England Biolabs, P8107S) per well. Conditions were incubated for 45 minutes at 85°C to lyse the cells. Polymerase chain reaction (PCR) was performed on each sample with Apex Taq RED 2X Master Mix (Genesee Scientific, 42-138B) using 1 µL of lysate and the primers in Table **S1**. PCR products were then run on a 2% agarose gel (Genesee Scientific, 20-102GP) at 110 V for 45 to 60 minutes and imaged using the ChemiDoc MP Imaging System (BioRad, 12003154).

#### Fluorescence imaging

Fluorescence images were taken on a Keyence All-in-One Fluorescence Microscope (Keyence, BZ-X800) at 10X magnification. Scale bars represent 100 µm.

### Quantification and statistical analysis

#### Analysis of flow cytometry data

Flow cytometry was performed on an Attune NxT Flow Cytometer (Thermo Fisher Scientific, A24858). See Table **S2** for corresponding fluorophores, lasers, and filters. Data were first analyzed in FlowJo (BD Biosciences, v10) to select cells using forward and side scatter gates (FSC-A, SSC-A) and single cells using forward scatter gates (FSC-A, FSC-H). The resulting single cell populations were analyzed in Python using the matplotlib (v3.10.3), numpy (v2.2.6), pandas (v2.2.3), rushd (v0.5.1), scipy (v1.15.3), and seaborn (v0.13.2) packages.

Fluorescence values are in arbitrary units plotted on a log scale and are shown as 1D or 2D kernel density estimates or as geometric means of gated populations. Cells were gated on expression of the downstream gene (for single-gene vectors) or on expression of both genes (for two-gene vectors), using gates defined by the 99.5^th^ percentile of untransduced cells or defined manually. Expression variability for the ComMAND vectors is shown as standard deviation of fluorescence (arbitrary units) of the gated population. Fold change comparing similar vectors with one different genetic part (vectors with a downstream EF1α promoter versus otherwise identical designs with a downstream EFS promoter, vectors with WPRE versus without WPRE) was computed by averaging the geometric means across biological replicates for the second vector, then dividing the geometric means for each biological replicate of the first vector by this value. Fold change comparing conditions treated with or without guanine was computed by dividing the geometric mean expression for the condition with guanine by the value for the condition without guanine within biological replicates.

Points represent means ±standard error across biological replicates. For transductions at a constant MOI, biological replicates are defined as independent transductions of separate batches of virus, for a total of *n* ≥ 3 for each vector. All statistics are two-sided Student’s t-tests. Annotations show fold changes computed by dividing the values for the indicated conditions averaged across biological replicates. For vectors with a Tet-ON inducible gene, the annotations always represent comparisons between conditions treated with dox.

#### Calculating functional titer

The fluorescence data from transduction of HEK293T cells with a two-fold serial dilution of virus was used to calculate a functional metric of viral titer. Specifically, transduction events were assumed to follow a Poisson process, where cells can be transduced by multiple viral particles and transduction events are independent. The expected number of transduction events per cell is the MOI, *λ*, meaning the probability of *k* events in a cell is

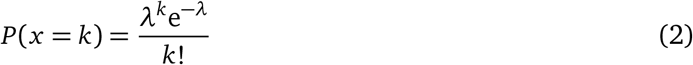

Across a population, this value is reflected by the fraction of cells with *k* transduction events. We can use the fraction of expressing cells (i.e., cells with at least one transduction event) to solve for MOI:

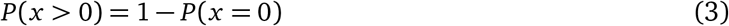

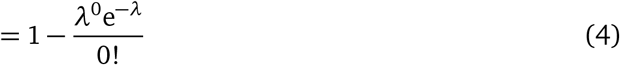

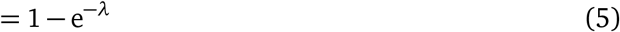

Combining Equations **1** and **5**, we have

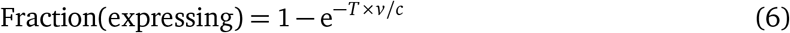

Using this equation, we fitted our data—the fraction of expressing cells as a function of virus volume (*v*) for six volumes of virus and a known number of cells (*c*)—to this equation for each batch of each vector to calculate titer (*T*, in TU/µL). Fraction expressing was calculated from expression gates defined by the 99.5^th^ percentile of untransduced cells. For curve fitting, we used the mean of two technical replicates per condition, and we excluded points where the fraction of expressing cells decreased for larger virus volumes. Titer values were calculated for untransduced cells (i.e., fraction expressing of 0.005 by definition) to serve as a reference. Normalized titer was computed relative to one vector per batch of virus to account for batch-to-batch variability. Titer is plotted on a log scale in TU/µL, or is unitless for normalized titer. Points represent means ±standard error across biological replicates. For titer calculations, biological replicates are defined as transductions of separate batches of virus, with *n* ≥ 3 batches per vector. All statistics are two-sided Student’s t-tests.

#### Calculating direct conversion yield

To calculate yield for direct conversion of mouse embryonic fibroblasts to induced motor neurons (iMNs), the iMN population was first defined as cells with high *Hb9*::GFP expression measured via fluorescence. The number of such cells was divided by the number of cells seeded at the beginning of the experiment, then normalized to the untreated condition (neither dox nor guanine added) for each lentiviral vector (i.e., vectors expressing HRAS^G12V^ with or without the splicing switch) for each biological replicate. Points represent means ±standard error across biological replicates. Biological replicates are defined as separate batches of donor cells in *n* = 3 independent conversion experiments, for a total of *n* = 5.

## Acknowledgments

We thank Shiva Razavi, Jonathan C. Chen, and James J. Collins at MIT for the splicing switch. A manuscript detailing the splicing switch, titled “A framework for programmable RNA-level gene control,” is currently in preparation. We thank Maria Castellanos, Mary Ehmann, Christopher Johnstone, Sneha Kabaria, Emma Peterman, and Nathan Wang for feedback on the manuscript.

## Funding

This work was supported by the National Institute of General Medical Sciences of the National Institutes of Health [R35-GM143033]; the National Science Foundation CAREER [2339986]; the Pershing Square Foundation MIND Prize; the National Science Foundation Graduate Research Fellowship [1745302 to K.S.L.]; and the MIT Health and Life Sciences Collaborative Graduate Fellowship [to K.S.L.].

## Author contributions

Following the CRediT taxonomy, K.S.L. contributed to conceptualization, methodology, investigation, validation, data curation, formal analysis, visualization, and writing (original draft, review & editing). B.A.L.-D. contributed to methodology, investigation, validation, formal analysis, and writing (review & editing). K.E.G. contributed to conceptualization, methodology, writing (review & editing), resources, supervision, funding acquisition, and project administration.

## Declaration of interests

There are no competing interests to declare.

## Notes

### Competing Interest Statement

The authors have declared no competing interest.

