## Supplement for "Engineering high-titer lentiviral vectors for robust expression of RNA-based gene circuits"

- Tables S1 and S2
- Figs. **S1**–**S7**

| Primer name | Sequence (5' to 3') |
|---|---|
| mCherry splicing switch forward | TGCAGGACGGCGAGTTCATC |
| mCherry splicing switch reverse | CGCGTTCGTACTGTTCCACGA |

**Table S1. Primers used for PCRs of genomic DNA from cells transduced with the splicing switch.** Primers bind in the mCherry coding sequence, flanking the site where the splicing switch is inserted.

| Gene | Fluorophore | Laser (nm) | Filter (nm) |
|---|---|---|---|
| Downstream gene (splicing switch vectors) | TagBFP | 405 | 440 / 50 |
| Downstream gene (other vectors)<br>iMN transgenic reporter | mGreenLantern<br>EGFP | 488 | 530 / 30 |
| Upstream gene (splicing switch vectors)<br>Upstream gene (other vectors) | mCherry<br>mRuby2 | 561 | 620 / 15 |

**Table S2. Channels and cytometer settings for flow cytometry experiments.**

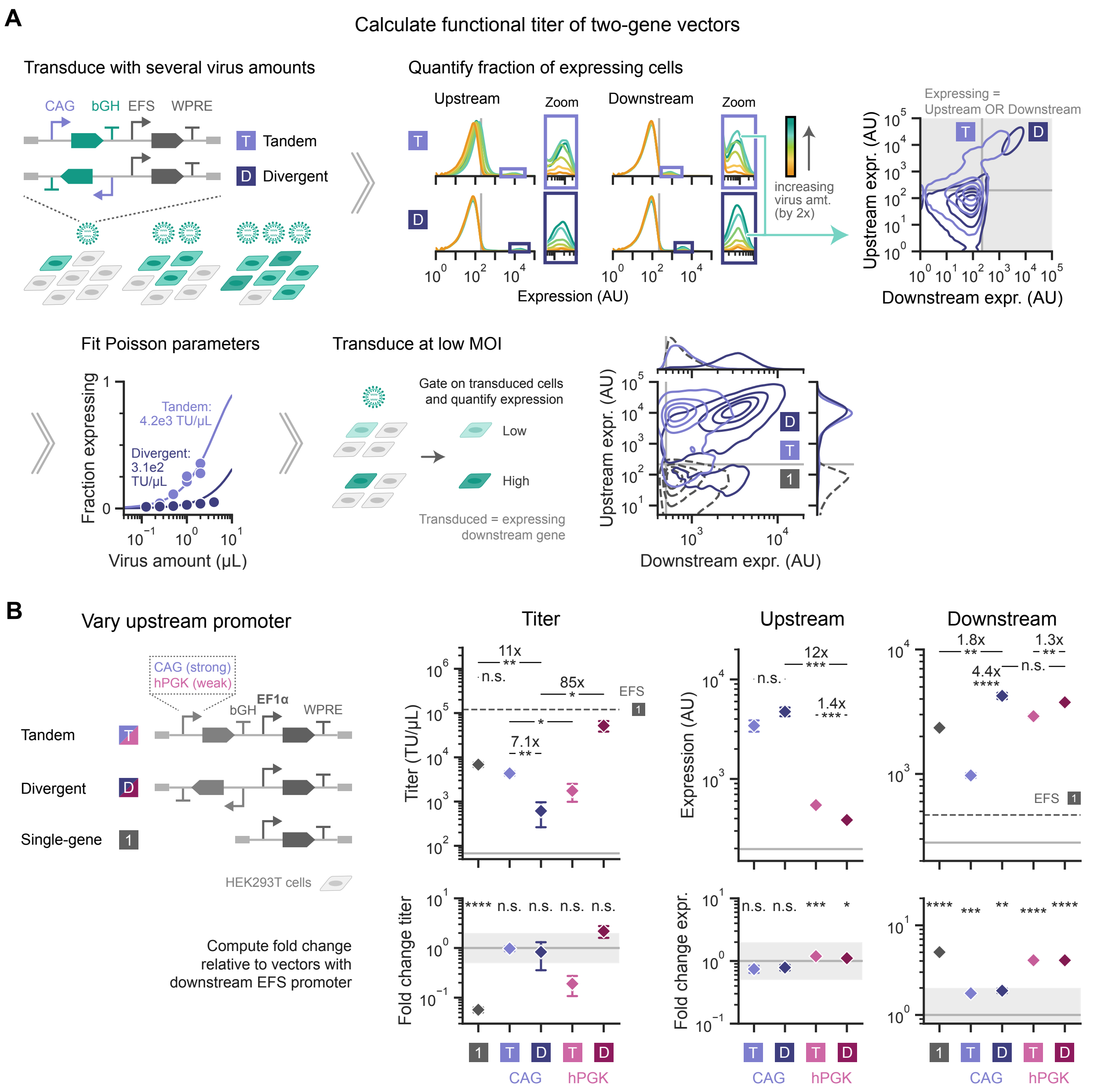


**Figure S1. Functional titer quantifies production efficiency for vectors with diverse genetic parts.**

**A.** Analogous to Fig. 1C, workflow for measuring functional titer and expression for two-gene vectors, shown for vectors with tandem (T) and divergent (D) syntax containing an upstream gene driven by the strong, synthetic CAG promoter and a second fluorescent protein driven by the EFS promoter downstream. **Top:** HEK293T cells were transduced with several different amounts of each virus. Expression of upstream and downstream genes were measured for each virus condition via fluorescence in arbitrary units (AU, log scale) from a flow cytometer. Light gray vertical line depicts the expression gate, used to calculate the fraction of expressing cells in each condition. Colored boxes indicate zoomed regions of the distributions. 2D density distributions depict expression (expr.) for conditions with the second-highest virus amount. Gray shading indicates cells considered expressing, i.e., above the expression gate for either gene. Plots show one representative batch of virus. **Bottom:** For the same vectors, fraction expressing as a function of virus volume was fit to a Poisson distribution to obtain titer in transducing units (TU) per μL virus. Points represent technical replicates for a representative batch of virus, and lines depict the Poisson curve fits. The computed titer is annotated on the plot, and virus volume is plotted on a log scale. To quantify cargo expression, fresh HEK293T cells were transduced at a low multiplicity of infection (MOI). 2D density distributions depict expression (AU, log scale) of cells expressing the downstream gene. A representative biological replicate is shown for the same two-gene vectors (solid lines) and for the EFS-driven single-gene vector from Fig. 1C (dashed lines).

*(continued)*

**Figure S1.** *(continued)*

**B.** Performance of single-gene and two-gene vectors as in Figs. 2C and 2D, except with EF1α expressing the downstream gene. **Top:** Titer is in transducing units (TU) per µL (log scale), and expression is the geometric mean fluorescence in arbitrary units (AU, log scale). Dark gray dashed lines show values for the single-gene EFS vector in Fig. 2C for reference. **Bottom:** Values above normalized to the analogous vectors with downstream EFS in Figs. 2C and 2D, plotted on a log scale. Solid gray line indicates a fold change of 1, and gray shading spans a two-fold change in either direction (0.5 to 2) for reference. Statistics compare unnormalized values.

Points represent means ± standard error for $n \geq 3$ biological replicates. Statistics are two-sided Student's t-tests, n.s. $p \geq 0.05$, * $p < 0.05$, ** $p < 0.01$, *** $p < 0.001$, **** $p < 0.0001$. Annotations show the fold change between indicated points.

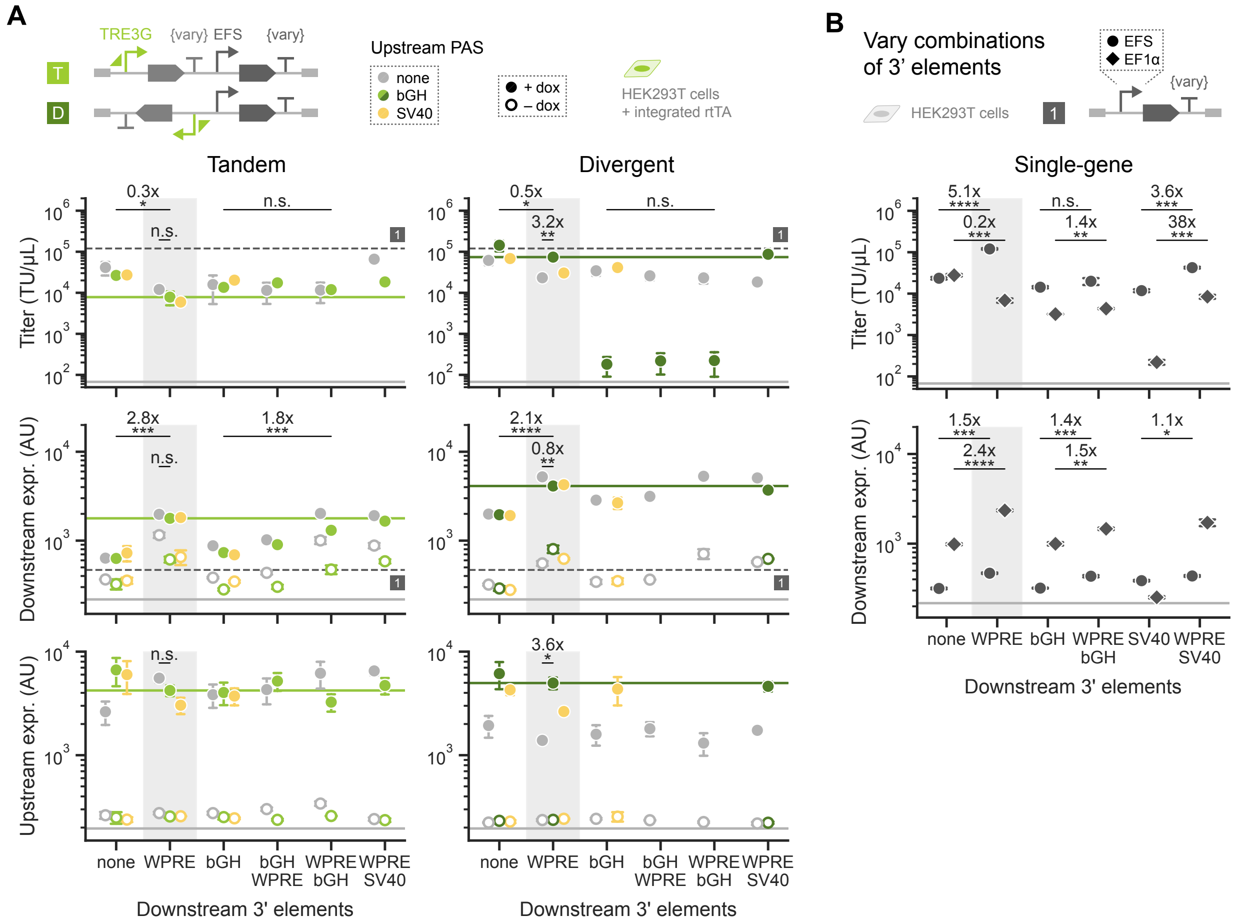


#### Figure S2. Varying downstream 3' elements and upstream PAS impacts vector titer and expression.

**A.** Performance of two-gene vectors with an inducible upstream gene as in Fig. 3, varying the upstream PAS and downstream 3' elements. Elements include combinations of the woodchuck hepatitis virus post-transcriptional regulatory element (WPRE), bovine growth hormone (bGH) polyadenylation signal (PAS), simian virus 40 (SV40) PAS, or none. Vectors were transduced into HEK293T cells containing a separately integrated rtTA cassette. Cells were treated with 1 µg/mL dox (filled points) or untreated (open points). Solid green lines depict values for the vectors in Fig. 3, and dark gray dashed lines show values for the single-gene vector in Fig. 2 for reference. Annotations show the fold change between indicated points for conditions treated with dox.

**B.** Performance of single-gene vectors with the EFS promoter (circles) or EF1α promoter (diamonds) and varying 3' elements. Vectors were transduced into HEK293T cells. Annotations show the fold change between indicated points.

Points represent means ± standard error for $n \geq 3$ biological replicates. Titer is in transducing units (TU) per µL (log scale). Expression is the geometric mean fluorescence in arbitrary units (AU, log scale). Solid light gray lines represent titers calculated for untransduced cells or expression gates. Gray shading highlights vectors with WPRE as the downstream 3' element, found in Figs. 2 and 3. Statistics are two-sided Student's t-tests, n.s. $p \geq 0.05$, * $p < 0.05$, ** $p < 0.01$, *** $p < 0.001$, **** $p < 0.0001$.

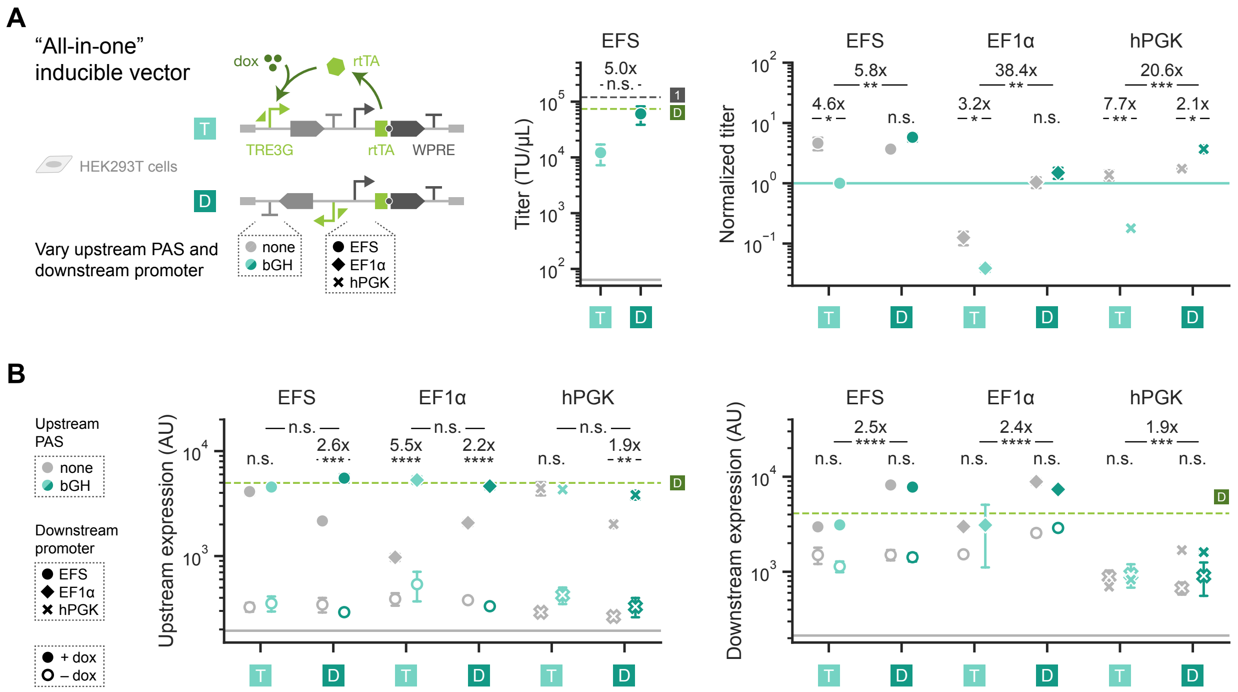


**Figure S3. "All-in-one" inducible vectors perform similarly with various genetic parts.**

**A.** "All-in-one" inducible vectors as in Fig. 4 were constructed varying the upstream PAS (none or bGH) and downstream promoter (EF1α, EFS, or hPGK). **Left:** Unnormalized viral titer for vectors in Fig. 4 in transducing units (TU) per μL (log scale). Dashed dark gray line depicts the value for the single-gene vector with the EFS promoter from Fig. 1C, and dashed green line shows the value for the inducible vector with divergent syntax from Fig. 3 for reference. Solid light gray line represents titer calculated for untransduced cells. **Right:** As in Fig. 4, viral titer is shown normalized to the vector with tandem syntax, upstream bGH PAS, and downstream EFS promoter (also indicated by the solid teal line) for each batch of virus (log scale).

**B.** Expression of the upstream and downstream genes for vectors in A transduced into HEK293T cells treated with 1 μg/mL dox (filled points) or untreated (open points). Solid light gray lines represent the expression gates. Dashed green lines depict values for the inducible vector with divergent syntax treated with dox from Fig. 3 for reference. Expression is the geometric mean fluorescence in arbitrary units (AU, log scale).

Points represent means ± standard error for $n \geq 3$ biological replicates. Statistics are two-sided Student's t-tests, n.s. $p \geq 0.05$, * $p < 0.05$, ** $p < 0.01$, *** $p < 0.001$, **** $p < 0.0001$. Annotations show the fold change between indicated points for conditions treated with dox.

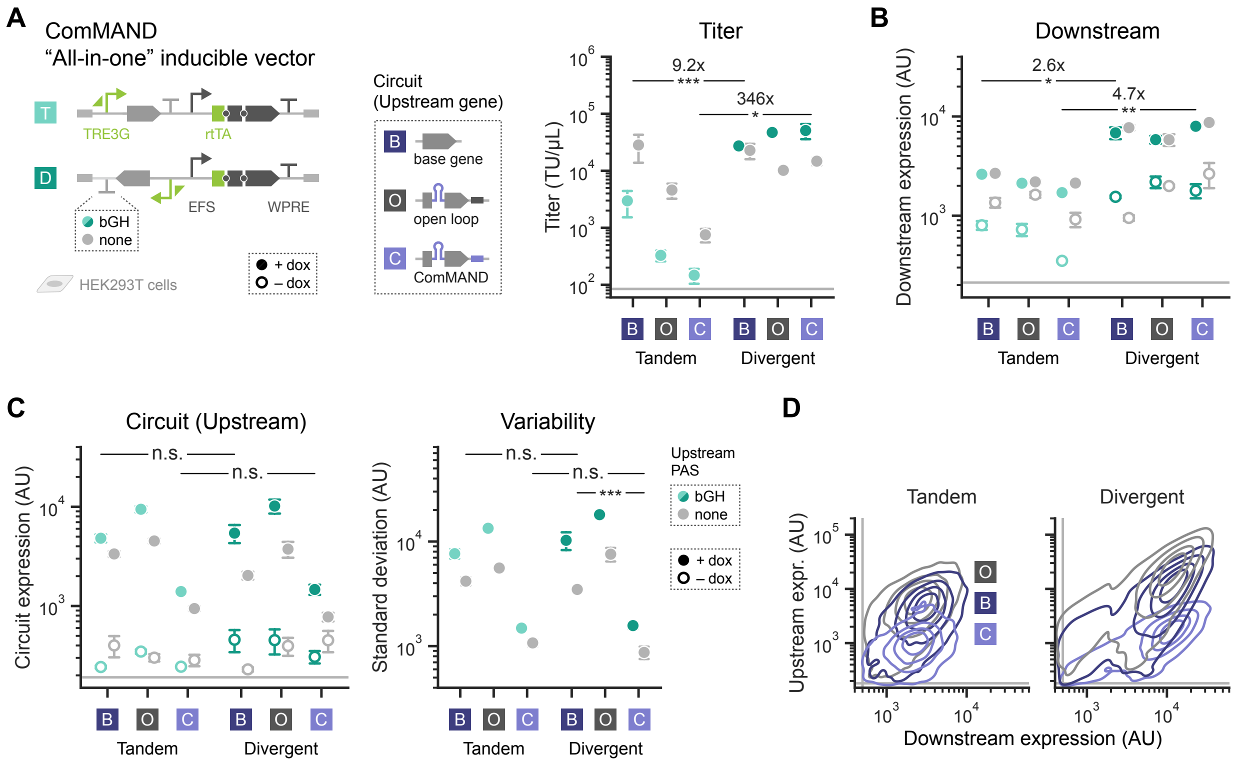


#### Figure S4. Additional "all-in-one" inducible vectors expressing ComMAND match trends.

**A.** "All-in-one" inducible vectors were constructed as in Fig. 5A varying the upstream PAS (bGH or none). The upstream gene expresses ComMAND (C), an open-loop circuit (O) containing the intronic microRNA and a mismatched target site as a control, or the output coding sequence alone (base gene, B). Here and in Fig. 5A, the downstream gene consists of rtTA, a fluorescent protein, and a puromycin resistance gene separated by 2A "self-cleaving" peptides. Titer is in transducing units (TU) per μL (log scale).

**B–C.** Expression of the downstream gene and the circuit (upstream gene) for HEK293T cells transduced with the same vectors and treated with 1 μg/mL dox (filled points) or untreated (open points). Expression is the geometric mean fluorescence in arbitrary units (AU, log scale) of populations gated on cells expressing both genes, and standard deviation (AU, log scale) is shown for expression of the upstream gene.

**D.** 2D density distributions depict upstream and downstream expression (AU, log scale) for a representative biological replicate of conditions in B–C, where tandem vectors lack a PAS on the upstream gene and divergent vectors include a bGH PAS as in Fig. 5A. Plots show cells treated with 1 μg/mL dox and gated on populations expressing both genes.

Points represent means ± standard error for $n \geq 3$ biological replicates. Solid light gray lines depict titer for untransduced cells or expression gates. Statistics are two-sided Student's t-tests, n.s. $p \geq 0.05$, * $p < 0.05$, ** $p < 0.01$, *** $p < 0.001$, **** $p < 0.0001$. Annotations show the fold change between indicated points for conditions treated with dox.

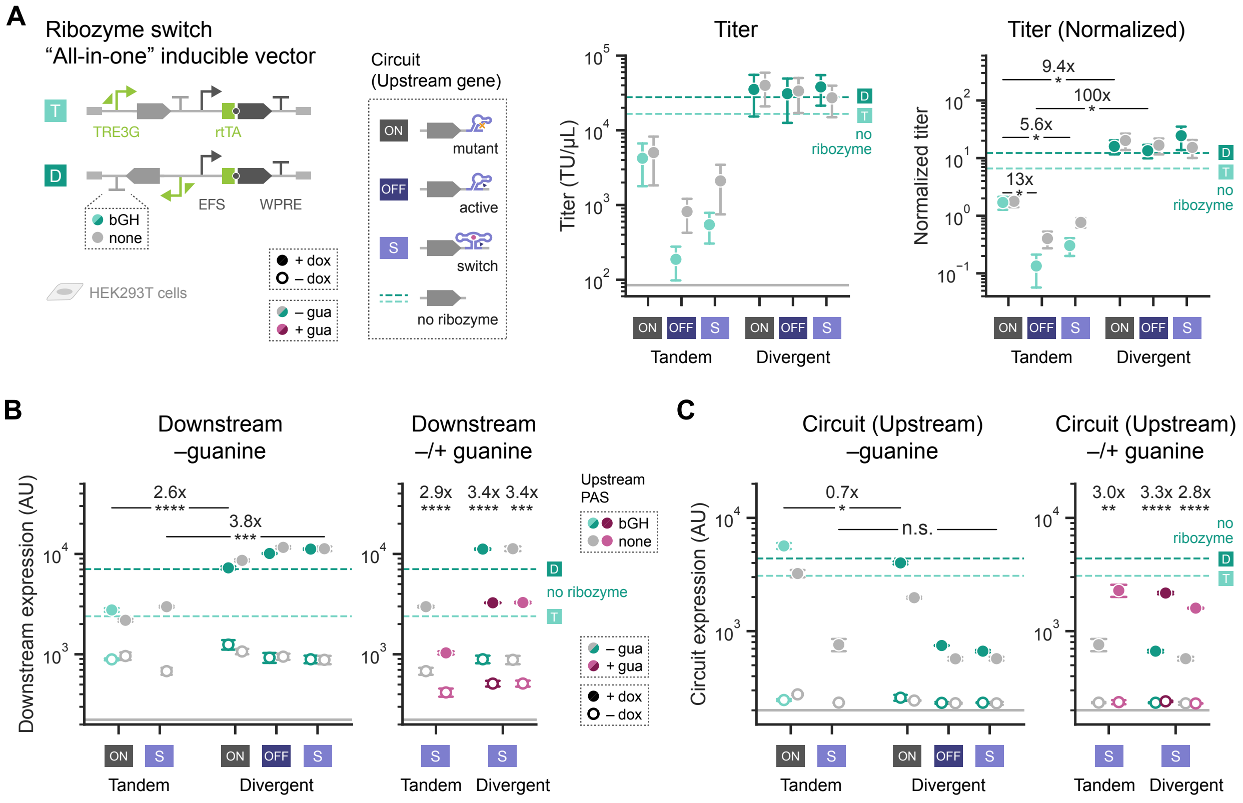


**Figure S5. Additional "all-in-one" inducible vectors expressing the ribozyme switch match trends.**

**A.** "All-in-one" inducible vectors were constructed as in Fig. **5B** varying the upstream PAS (bGH or none). The 3' UTR of the upstream gene contains either a mutant, minimally cleaving ribozyme (expression ON); a constitutively active ribozyme (expression OFF); a guanine (gua)-responsive ribozyme switch (S); or no ribozyme (dashed teal lines). **Left:** Titer is in transducing units (TU) per μL (log scale). **Right:** Titer is normalized to a vector lacking a ribozyme (not shown) for each batch of virus (log scale).

**B–C.** Expression of the downstream gene and the circuit (upstream gene) for HEK293T cells transduced with the same vectors and treated with 1 μg/mL dox (filled points) or untreated (open points). Vectors containing the ribozyme switch include additional conditions where cells were also treated with 100 μM guanine (pink), with or without dox. Expression is the geometric mean fluorescence in arbitrary units (AU, log scale).

Points represent means ± standard error for $n \geq 3$ biological replicates. Solid light gray lines depict titer for untransduced cells or expression gates. Statistics are two-sided Student's t-tests, n.s. $p \geq 0.05$, * $p < 0.05$, ** $p < 0.01$, *** $p < 0.001$, **** $p < 0.0001$. Annotations show the fold change between indicated points for conditions treated with dox.

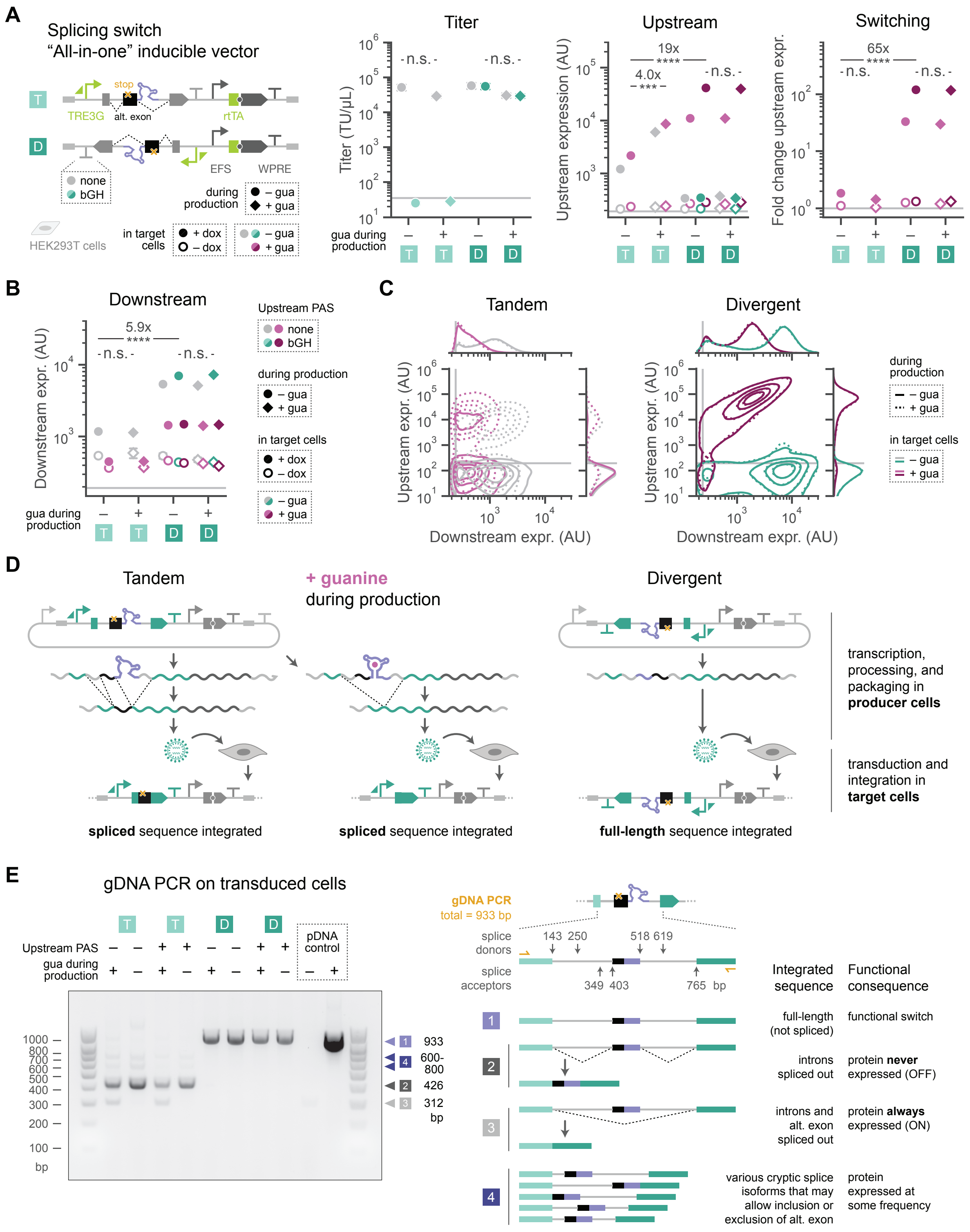


**Figure S6. Additional "all-in-one" inducible vector designs expressing a splicing switch.**

*(caption on next page)*

**Figure S6.** *(continued)*

**A–B.** "All-in-one" inducible vectors as in Fig. **6A** encode the splicing switch within the upstream gene, with divergent or tandem syntax and with no PAS or the bGH PAS on the upstream gene. Vectors were produced in HEK293T cells cultured without (circles) or with (diamonds) 100 µM guanine (gua). Titer is in transducing units (TU) per µL (log scale). Expression of upstream and downstream genes is the geometric mean fluorescence in arbitrary units (AU, log scale) for HEK293T cells treated with (filled points) or without (open points) 1 µg/mL dox and with (pink) or without 100 µM guanine. Fold change in upstream expression shows conditions treated with guanine relative to those without.

**C.** 2D density distributions depict upstream and downstream expression for the conditions in A–B.

**D.** The splicing switch can interfere with virus production. When encoded on the sense strand of the viral genome, the switch may retain activity in producer cells, leading to packaging and integration of spliced sequences. Producer cells cultured with or without guanine can generate different splice products for tandem vectors, resulting in integration of different sequences lacking the splicing switch in target cells (undesired). In contrast, for the divergent vector, the viral transcript is not spliced in producer cells, leading to integration of the full-length sequence in target cells (desired).

**E. Left:** The gel from Fig. **6B** is shown uncropped. The gel depicts PCR products amplified from genomic DNA (gDNA) of cells transduced with the vectors in A. Vectors have the bGH PAS (+) or no PAS (–) on the upstream gene and were produced in the presence (+) or absence (–) of 100 µM guanine. As controls, plasmid DNA for one of the vectors was included in a separate reaction (pDNA +), or no template was added to the reaction (pDNA –). Lengths of the ladder bands are labeled in base pairs (bp), and the most prominent bands in the conditions are annotated with their predicted lengths and sequence. **Right:** The amplicon spans the site where the splicing switch was added, and primers (orange) bind in the mCherry coding sequence. Predicted splice donor and acceptor sites are annotated (arrows) with their location on the amplicon (in bp). Predicted splice products and integrated sequences are shown for the most prominent bands on the gel, noting their functional consequence for protein expression in target cells.

Solid light gray lines depict titer for untransduced cells or expression gates. Points represent means ± standard error for $n \geq 3$ biological replicates. Statistics are two-sided Student's t-tests, n.s. $p \geq 0.05$, * $p < 0.05$, ** $p < 0.01$, *** $p < 0.001$, **** $p < 0.0001$. Annotations show the fold change between indicated points for conditions treated with dox. Alt. exon, alternative exon.

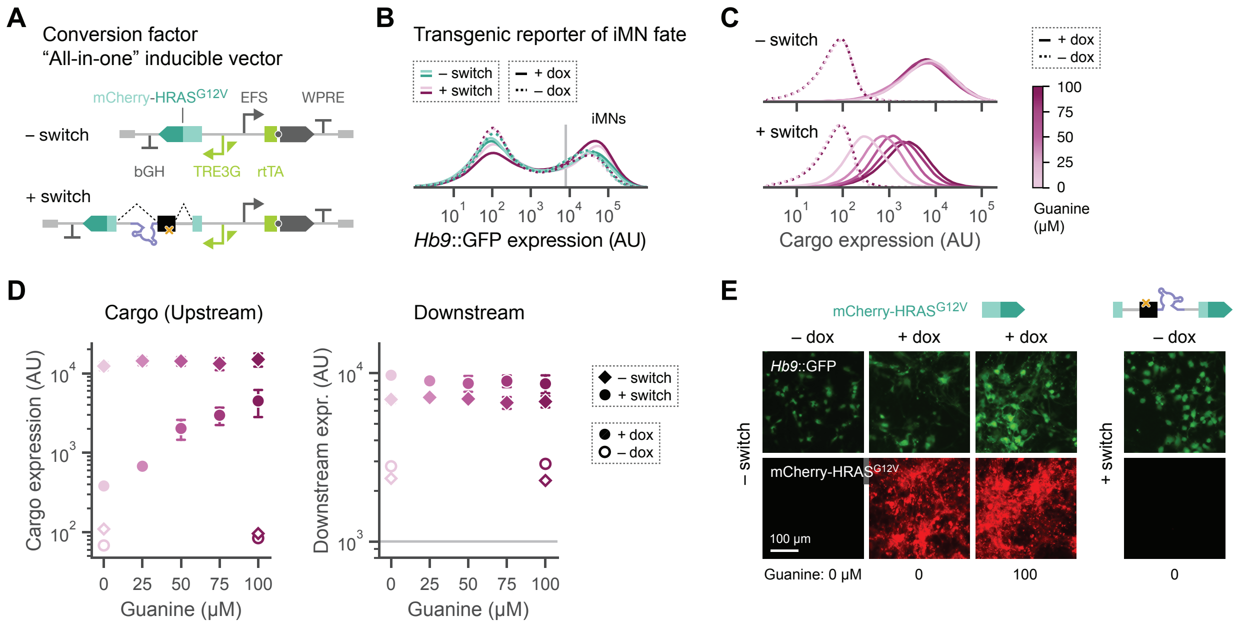


### Figure S7. A splicing switch can regulate expression of a conversion factor during reprogramming.

**A.** The splicing switch regulates expression of the oncogenic HRAS mutant HRAS$^{G12V}$, which is fused to the fluorescent protein mCherry and delivered via the the best-performing divergent "all-in-one" inducible vector. Full vectors with or without the splicing switch used in Figs. **6C** and **6D** are shown.

**B.** Primary mouse embryonic fibroblasts were directly converted to induced motor neurons (iMNs) by virally delivering conversion factors including the vectors in A containing (pink) or lacking (teal) the splicing switch. Cells were treated with 1 μg/mL dox (solid line) or untreated (dashed line) for the duration of the conversion process. The transgenic reporter *Hb9::GFP* is an indicator of iMN fate. At 14 days post-transduction, GFP expression was quantified via fluorescence in arbitrary units (AU, log scale) to identify iMNs. Solid light gray line indicates the expression gate.

**C.** Expression of the mCherry-HRAS$^{G12V}$ cargo (upstream gene) is measured via fluorescence (AU, log scale) for iMNs identified as in B. Plots show distributions for cells transduced with vectors containing or lacking the splicing switch, treated with 1 μg/mL dox (solid lines) or without dox (dashed lines), and treated with 0–100 μM guanine.

**D.** Cargo and downstream expression are shown for conditions in C as a function of guanine concentration. Expression is geometric mean fluorescence (AU, log scale). Solid light gray line shows the expression gate. Points represent means ± standard error for $n = 5$ biological replicates.

**E.** As in Fig. **6D**, images show fluorescence of *Hb9*::GFP (top) and mCherry-HRAS$^{G12V}$ (bottom) at 14 days post-transduction for cells transduced with vectors lacking or containing the splicing switch, treated with or without dox, and treated with varying concentrations of guanine. Scale bar represents 100 μm.
